# PTGER2-β-Catenin Axis Links High Salt Environments to Autoimmunity by Balancing IFNγ and IL-10 in FoxP3^+^ Regulatory T cells

**DOI:** 10.1101/379453

**Authors:** Tomokazu Sumida, Matthew R. Lincoln, Chinonso M. Ukeje, Donald M. Rodriguez, Hiroshi Akazawa, Tetsuo Noda, Atsuhiko T. Naito, Issei Komuro, Margarita Dominguez-Villar, David A. Hafler

## Abstract

Foxp3^+^ regulatory T cells (Tregs) are the central component of peripheral immune tolerance. While dysregulation of the Treg cytokine signature has been observed in autoimmune diseases such as multiple sclerosis (MS) and type 1 diabetes, the regulatory mechanisms balancing pro- and anti-inflammatory cytokine production are not known. Here, we identify imbalance between IFNγ and IL-10 as a shared Treg signature, present in patients with MS and under high salt conditions. By performing RNA-seq analysis on human Treg subpopulations, we identify β-catenin as a key regulator that controls the expression of IFNγ and IL-10. The activated β-catenin signature is enriched specifically in IFNγ^+^Tregs in humans, and this was confirmed *in vivo* with Treg-specific β-catenin-stabilized mice exhibiting lethal autoimmunity with a dysfunctional, IFNγ-producing, Treg phenotype. Moreover, we identify PTGER2 as a major factor balancing IFNγ and IL-10 production in the context of a high salt environment, with skewed activation of the β-catenin/SGK1/Foxo axis in IFNγ^+^Tregs. These findings identify a novel molecular mechanism underlying inflammatory Tregs in human autoimmune disease and reveal a new role for a PTGER2-β-catenin loop in Tregs linking environmental high salt conditions to autoimmunity.

## Introduction

The homeostatic maintenance of T cells is finely tuned by Foxp3^+^ regulatory T cells (Tregs). Tregs play a distinct role from the other CD4^+^ T cells in dampening prolonged inflammation and preventing aberrant autoimmunity^1^. Foxp3 has been shown to be a unique master regulator of Treg function and differentiation, and loss of function of Foxp3 is known to cause severe autoimmune and inflammatory diseases, such as immunodysregulation polyendocrinopathy enteropathy X-linked (IPEX) syndrome in humans and the *scurfy* phenotype in mice^2, 3^. Although Tregs are potent suppressors of immune function, the number of Tregs is often normal in a variety of autoimmune diseases, including multiple sclerosis (MS), inflammatory bowel disease (IBD), and type 1 diabetes (T1D)^4, 5^. These observations suggest that not only a quantitative, but also a functional dysregulation of Tregs contributes to the development of autoimmunity.

Tregs display their suppressive capacity in peripheral immune responses through both contact-dependent and cytokine-mediated mechanisms^6^. It has been shown that Tregs demonstrate substantial heterogeneity and that the balance between pro- and anti-inflammatory populations is finely regulated to maintain immunologic homeostasis^7^. Recent studies provided evidence that IFNγ identifies dysfunctional Tregs in patients with autoimmunity (MS^8^ and T1D^9^) and cancer (glioblastoma^10^). Additionally, Tregs producing the anti-inflammatory cytokine IL-10 have been reported to play prominent roles in suppressing the immune response at environmental interfaces^11^ and development of mature memory CD8^+^ T cells to prevent autoimmunity and chronic infection in mice^12^. These studies suggest that the balance between IFNγ and IL-10 production in Tregs may be central in the maintenance of immune homeostasis; however, the molecular mechanisms underlying this regulatory balance are not known.

Human autoimmune disease results from an interplay between genetic factors and environmental triggers. In this regard, MS is an autoimmune disease that results from the complex interaction of predominantly common genetic variants and environmental factors^13^, with 233 common risk haplotypes identified to date^14,15^. Several environmental factors are associated with an increased risk of MS including vitamin D insufficiency, smoking, obesity, and a high salt diet^16^. Increased dietary salt intake has recently been associated with increased clinical disease activity in MS patients. Previously, we have observed that a high salt diet (HSD) exacerbated neuroinflammation in the experimental autoimmune encephalomyelitis (EAE) model of MS^17, 18^, and that higher salt concentration within the physiological range skewed naïve CD4^+^ T cells into pro-inflammatory Th17 cells^17, 18^ and impaired Treg suppressive function through induction of IFNγ expression^19^. Studies using murine models of autoimmune disease are accumulating to support this theory^20, 21^ and recent magnetic resonance imaging (MRI) studies investigating the association between interstitial sodium content and inflammatory diseases in humans revealed higher sodium accumulation in acute MS lesions compared to chronic lesions, suggesting higher sodium concentration within the pathogenic microenvironment in MS brain^22^. However, it remains unknown whether a high salt diet has a direct impact on MS clinical activity^23^.

β-catenin is an essential component of the canonical Wnt signaling pathway and is known to be involved in a variety of biological processes including carcinogenesis, stem cell maintenance, organogenesis, and aging^24, 25^. In the absence of Wnt ligands, β-catenin is actively phosphorylated by a destruction complex composed of glycogen synthase kinase 3β (GSK3β), axis inhibition protein (Axin), and casein kinase 1 (CK1), and the complex is subsequently degraded by the ubiquitin-proteasome system. Binding of Wnt ligands to their receptors inactivates the destruction complex, leading to the accumulation of unphosphorylated β-catenin protein, which can interact with TCF/LEF transcription factors^26^. While β-catenin and canonical Wnt signaling have been studied in the context of memory CD8^+^ T cell development, T helper cell differentiation, and Treg function^27, 28, 29, 30^, results differ from study to study, depending on the experimental model utilized. In Tregs, it was first demonstrated that forced induction of constitutively active β-catenin in human Tregs leads to induction of survival gene expression and promotion of anti-inflammatory function^28^. In contrast, recent work suggested that pharmacological activation of Wnt signaling modulates regulatory activity of Foxp3 and disrupts Treg function^29^. This result was partially supported by another study using transgenic mice in which β-catenin was stabilized specifically in CD4^+^ T cells^30, 31^. Although these two recent studies suggested β-catenin as a driving factor of Treg dysfunction, the specific mechanisms by which β-catenin affects Treg function and their role in modulating cytokine production by Tregs, in particular in the context of human autoimmune disease, is poorly understood.

Here, we show that the imbalance between IFNγ and IL-10 is a shared Treg signature observed in the patients with multiple sclerosis and high salt environment. By performing unbiased RNA-seq analysis on human Treg subpopulations, we dissect Treg heterogeneity and identify β-catenin as central in maintaining Treg function and regulating both IFNγ and IL-10 cytokine production. Moreover, we clarify a previously unknown role for β-catenin in mediating the high salt-induced pro-inflammatory signature by creating a feed forward loop with PTGER2, which is uniquely upregulated under high salt conditions. Finally, we demonstrate the clear association between IFNγ, β-catenin, and PTGER2 expression in Tregs from MS patients. Our findings suggest that the β-catenin-PTGER2 axis serves as a bridge between environmental factors and autoimmune disease by modulating Treg function and this axis may be involved in the pathogenesis of autoimmune disease.

## Results

### Treg cytokine imbalance in multiple sclerosis and high salt environment

We and others have previously identified a pro-inflammatory Treg population characterized by the secretion of IFNγ. This population is dysfunctional both *in vitro* and *in vivo*, and a high frequency of this population is associated with autoimmune disease and cancer^8, 9, 10^. However, the balance between pro- and anti-inflammatory Treg populations has not been defined. To address this question, we evaluated the production of pro-inflammatory (IFNγ) and anti-inflammatory (IL-10) cytokines by circulating human Tregs from healthy subjects and patients with MS by flow cytometry. Based on our observation that CD25^hi^CD127^low-neg^CD45RO^+^ Tregs (memory Tregs; mTregs) are the major source for effector cytokine expression in human Tregs (Supplementary Fig. 1), we focused on mTregs, so as to avoid the potential bias caused by the variable ratio of naïve Tregs and memory Tregs between subjects. We found that mTregs isolated from MS patients (MS-Tregs) produced more IFNγ and less IL-10 compared to healthy donors, and the ratio of IFNγ to IL-10 producing Tregs further highlights this imbalance (Fig. 1a, b). Furthermore, we examined the mRNA expression of *IFNG* and *IL10* genes in mTregs without PMA/iomomycin stimulation, better reflecting the situation *in vivo*, and identified a trend similar to that seen in protein expression (Fig. 1c).

**Figure 1.**
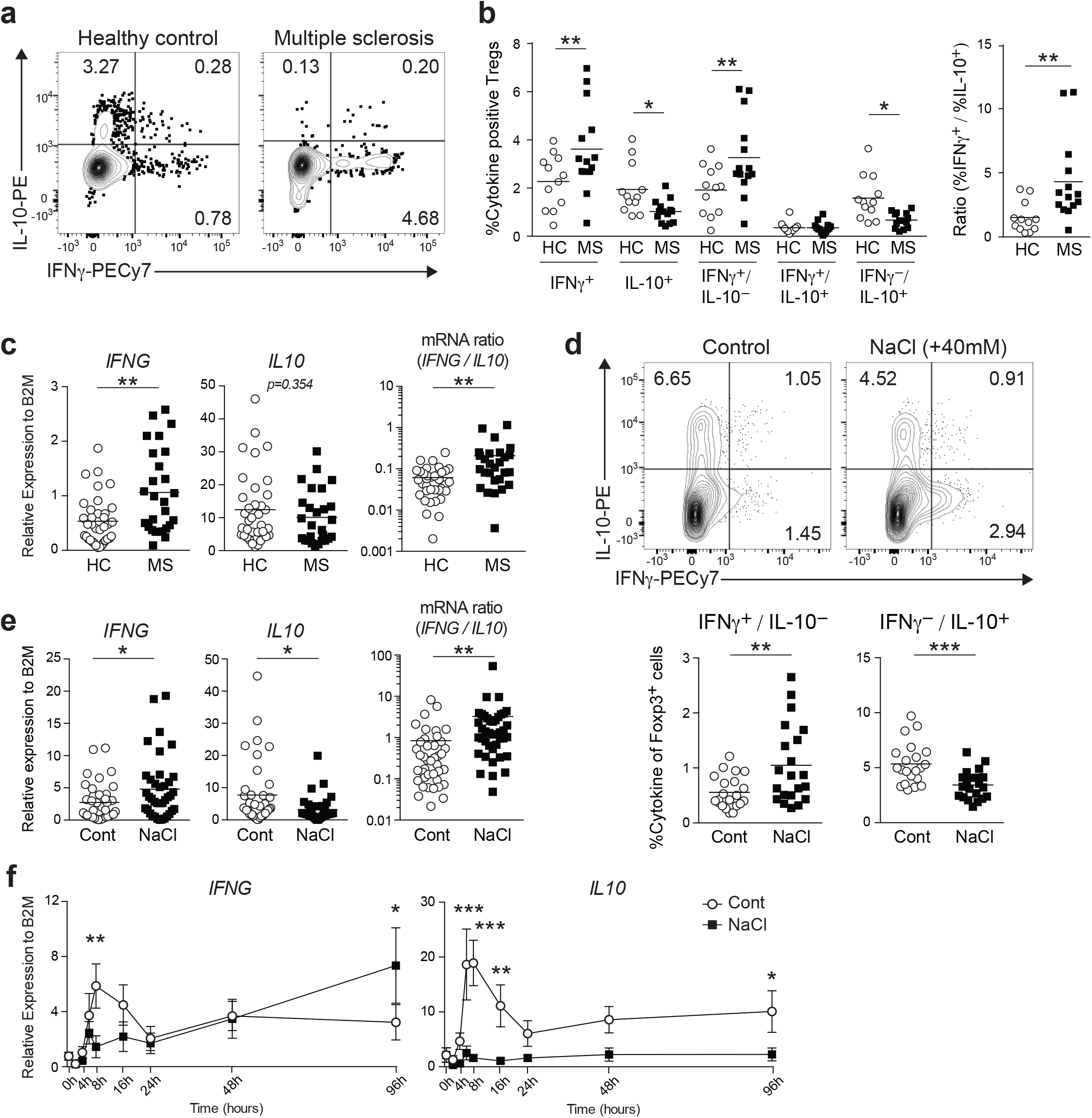
IFNγ/IL-10 balance of human Tregs in MS and high salt treatment. **(a)** Representative flow cytometric analysis of *ex vivo* human Tregs (CD4^+^CD25^hi^CD127^neg^CD45RO^+^) isolated from healthy donor and MS patient. FACS isolated Tregs were stimulated with PMA/iomomycin for 4 h followed by intracellular staining for IFNγ and IL-10. **(b)** IFNγ and IL-10 cytokine profiles of *ex vivo* human Tregs. Percentage of IFNγ and/or IL-10 producing Tregs was shown (left). Ratio between IFNγ positive and IL-10 positive Tregs was plotted (right). (HC; n=12, MS; n=14) \**P*<0.05, \*\**P*<0.01 (two-way ANOVA with Sidak’s multiple comparisons test). **(c)** *IFNG* and *IL10* gene expression was determined on *ex vivo* Tregs by qPCR. Ratio between *IFNG* and *IL10* expression was shown at the right (HC; n=36, MS; n=27). \*\**P*<0.01 (two-tailed unpaired Student’s *t*-test). **(d)** Representative flow cytometric analysis of IFNγ and IL-10 production in human Tregs stimulated with anti-CD3 and anti-CD28 in the normal media (Control) or media supplemented with additional 40 mM NaCl (NaCl) for 96 h. Percentage of IFNγ and/or IL-10 producing Tregs was shown at the right (n=21 subjects). **(e)** *IFNG* and *IL10* mRNA expression was assessed at 96 h following stimulation as in **(d)** (n=41 subjects). Ratio between *IFNG* and *IL10* expression was plotted (right). \**P*<0.05, \*\**P*<0.01 (two-tailed unpaired Student’s *t*-test). **(f)** mRNA expression kinetics of *IFNG* and *IL10* from 9 different time points were plotted. Data were represented as mean +/− SEM (n=6 subjects). \**P*<0.05, \*\**P*<0.01, \*\*\**P*<0.001 (two-way ANOVA with Sidak’s multiple comparisons test).

We recently demonstrated that Tregs exposed to high salt concentrations exhibited a dysfunctional phenotype with a pro-inflammatory cytokine signature skewed towards IFNγ^19^. We sought to determine whether high salt could also impair the IFNγ/IL-10 balance and found that the high salt environment caused an increase in IFNγ and decrease in IL-10 production in human mTregs after 96 h in culture (Fig. 1d, e). Gene expression kinetics of *IFNG* and *IL10* by using qPCR identified early (8 h) and late (96 h) waves of gene expression. High salt stimulation suppressed the early wave of *IFNG* and *IL10*, and enhanced the late wave of *IFNG* but not *IL10* (Fig. 1f). These findings suggest that the imbalance of IFNγ/IL-10 induced by continuous exposure to high salt conditions, which is not observed at the phase of acute response to high salt stress, might capture the dysfunctional Treg properties in the setting of autoimmunity.

### β-catenin as a regulatory factor for IFNγ/IL-10 production in human Tregs

While our previous investigations have identified the AKT pathway as important in inducing IFNγ secretion with loss of Treg function^32^, the molecular mechanisms underpinning the balance between IFNγ and IL-10 in Tregs are largely unknown. To address this question, we performed RNA-sequencing (RNA-seq)-based genome-wide transcriptome analysis on human Treg subsets defined by IFNγ and IL-10 production. mTregs isolated from peripheral mononuclear cells of healthy subjects were stimulated with PMA/iomomycin for 4 h *ex vivo*. After applying cytokine capture kits for IFNγ and IL-10, we sorted four different subpopulations (IFNγ single positive (IFNγSP), IL-10 single positive (IL10SP), IFNγ and IL-10 double positive (DP), and double negative (DN)), and we performed RNA-seq on each subpopulation (Fig 2a). We identified 672 differentially expressed genes between IFNγSP and IL10SP and the four populations could be distinguished by their gene expression profiles (Fig. 2b). Of note, the IFNγ-producing populations were highly distinct from IFNγ-negative populations, suggesting that IFNγ-secreting Tregs represent a more dominant signature than IL-10-secreting Tregs. We also identified ten clusters of co-expressed genes (C1-C10) across the populations. IFNγ and IL-10-associated genes are enriched in C9/C10 (e.g. *CXCR3, CD226*, and *NKG7*) and C1/C2 (e.g. *MAF, SOCS3*, and *NOTCH2*), respectively.

**Figure 2.**
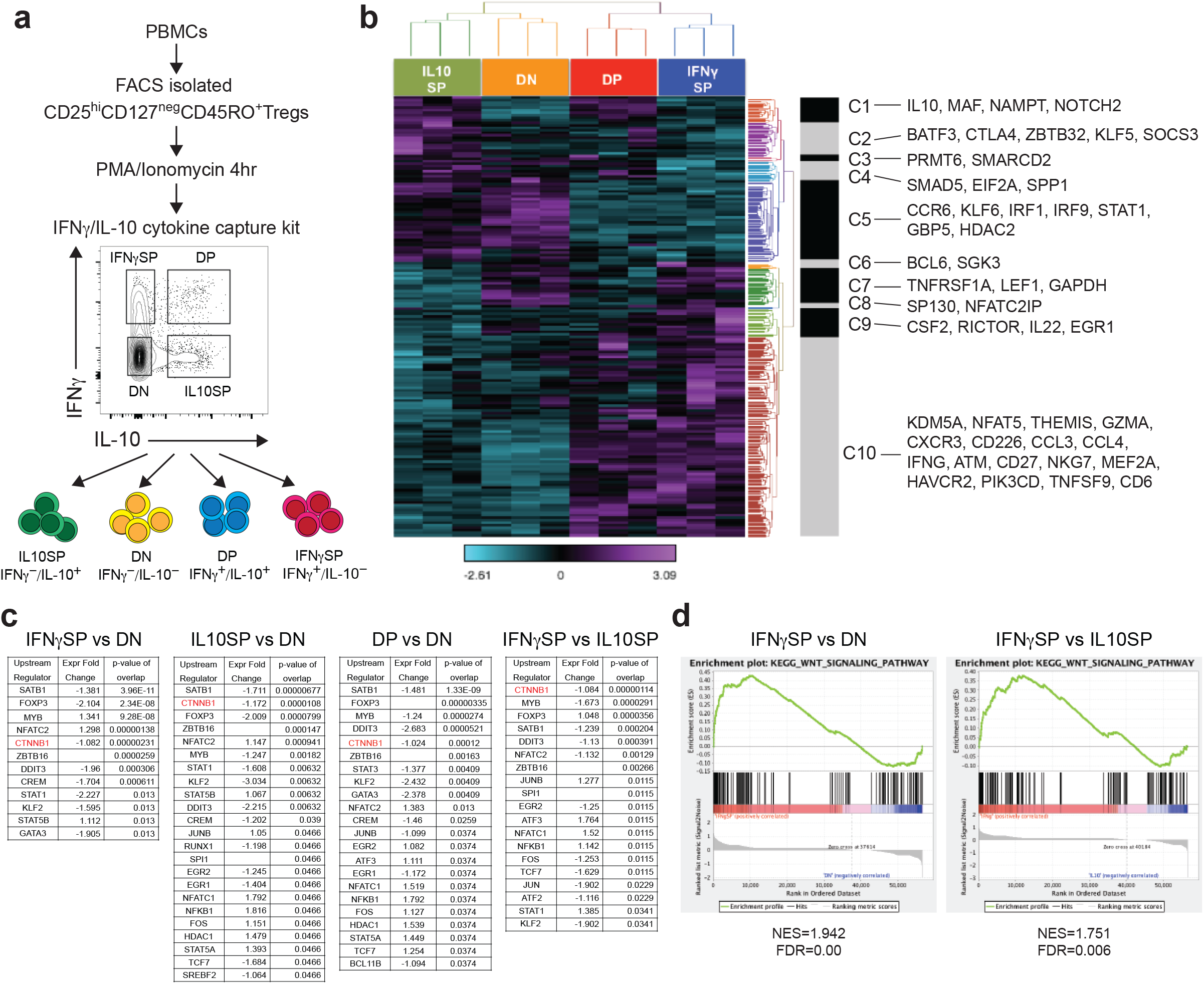
Transcriptional profiling of IFNγ/IL-10 producing human Treg subsets. **(a)** Experimental workflow for RNA-seq with IFNγ/IL10 producing Treg subpopulations. **(b)** Heatmap of 672 differentially expressed genes between IFNγSP and IL10SP. 10 clusters are identified and representative genes for each cluster are shown. **(c)** Lists of the top-ranking genes identified by IPA analysis as upstream regulators between each Treg subpopulations. Tables show statistically significant (overlap *P* value <0.05) upstream regulators in each comparison (Genes that could not be calculated for fold change were blank). *CTNNB1* gene, which codes β-catenin protein, was highlighted in red. **(d)** GSEA enrichment plots of KEGG Wnt signaling pathway between INFγSP vs. DN and INFγSP vs. IL10SP. Normalized enrichment score (NES) and false discovery rate (FDR) were represented at the bottom of each plot.

To predict the key transcriptional regulators that account for IFNγ and IL-10 production in Tregs, we performed an upstream regulator analysis in Ingenuity Pathway Analysis (IPA), using differentially expressed genes from each population (Fig. 2c). We identified β-catenin (*CTNNB1*) as one of the top upstream regulators in the Treg populations producing IFNγ and/or IL-10 compared to DN. Intriguingly, β-catenin was ranked as the top-ranked upstream regulator in the comparison between IFNγSP and IL10SP. Together, these results suggest that β-catenin plays a critical role in driving the production of both IFNγ and IL-10 in Tregs, especially for IFNγ. We also identified several upstream regulators that have been demonstrated to have critical roles in maintaining Treg function, including *MYB*^33^, *SATB1*^34^, *NFATC2*^35^, *and KLF2*^36^, suggesting that our upstream regulator analysis provides a reliable readout.

In agreement with these findings, gene-set-enrichment analysis (GSEA) applied to transcriptional profiles in each population identified significant enrichment of the Wnt/β-catenin signaling pathway in IFNγ-producing Treg subsets (Fig. 2d, Supplementary Fig. 2a). Notably, IFNγSP exhibited the highest enrichment score for the Wnt/β-catenin signaling pathway. Further GSEA analysis with different gene sets also provided similar results (Supplementary Fig. 2b). Taken together, these findings suggest that Wnt/β-catenin signaling is more activated in IFNγ-secreting Tregs than in other subpopulations of circulating human Tregs.

### β-catenin is stabilized in the IFNγ secreting Treg population

As β-catenin has multiple cellular functions and interacts with a variety of regulatory molecules, there is a growing body of evidence indicating its potential role in multiple pathological conditions, including MS^37, 38^. We first confirmed that β-catenin is stabilized and transcriptionally active in IFNγSP compared to DN in *ex vivo* Tregs by examining the level of active β-catenin (ABC), the dephosphorylated form of β-catenin with established active transcriptional activity^39^ (Fig. 3a). Notably, the DP and IL10SP also exhibited increased active β-catenin expression compared to DN *ex vivo*, suggesting that β-catenin signaling is important not only for IFNγ but also for IL-10 production in Tregs, consistent with our upstream regulator and enrichment analyses (Fig. 2c, Supplementary Fig. 2a). To exclude the possibility that PMA/iomomycin stimulation affected β-catenin stability, we also measured active β-catenin levels in CXCR3^+^ Th1-like Tregs, which contain most of the IFNγ-producing Tregs^40^ without PMA/iomomycin stimulation; these analyses confirmed that active β-catenin expression was significantly increased in the CXCR3^+^Th1-like Treg population (Fig. 3b). In agreement with these data, the downstream β-catenin target genes, *AXIN2* and *TCF7*, and the protein TCF-1 (encoded by *TCF7*) were upregulated in IFNγSP compared to DN *ex vivo* (Fig. 3c, Supplementary Fig. 3c). This was consistent with our previously published microarray data for IFNγ-positive and IFNγ-negative Tregs^32^ (Supplementary Fig. 3a).

**Figure 3.**
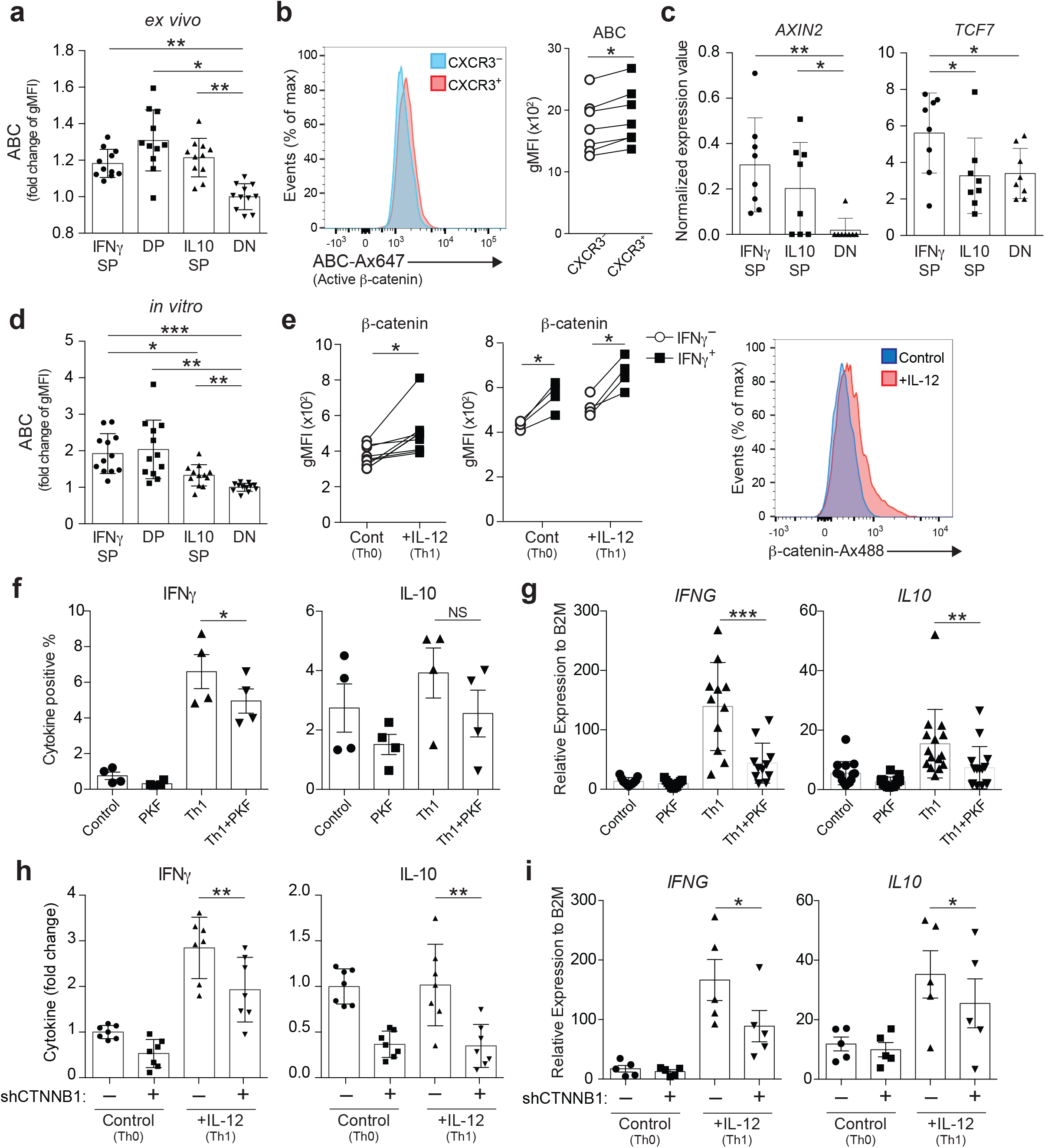
β-catenin is stabilized in the IFNγ secreting Treg population. **(a)** Relative expression level of Active β-catenin (ABC) on *ex vivo* Treg subpopulations analyzed by flow cytometry (n=11 subjects). Fold change in gMFI over DN were depicted. \**P*<0.05, \*\**P*<0.01 (one-way ANOVA with Tukey’s multiple comparisons test). gMFI, geometric mean fluorescence intensity. **(b)** Expression level of ABC between CXCR3^−^ and CXCR3^+^ *ex vivo* Tregs from healthy controls. Representative histogram (left) and summary of results (n=7 subjects) (right). \**P*<0.05 (two-tailed paired Student’s *t*-test). **(c)** Gene expression of Wnt/ β-catenin signaling target genes (*AXIN2* and *TCF7*) assessed by RNA-seq. \**P*<0.05, \*\**P*<0.01 (one-way ANOVA with Tukey’s multiple comparisons test). **(d)** Relative expression level of ABC on Tregs stimulated with anti-CD3 and anti-CD28 for 4 days, followed by 4 h PMA/iomomycin stimulation and intracellular cytokine staining for IFNγ and IL-10 (n=12 subjects). Fold change in gMFI over DN were depicted. \**P*<0.05, \*\**P*<0.01, \*\*\**P*<0.001 (one-way ANOVA with Tukey’s multiple comparisons test). **(e)** Expression level of β-catenin on Tregs stimulated with anti-CD3 and anti-CD28 in the presence (Th1) or absence (Th0) of IL-12 for 4 days. β-catenin level was determined directly by intracellular staining (left) (n=9 subjects) and β-catenin level on IFNγ positive/negative Treg populations was determined after 4 h PMA/iomomycin stimulation (middle) (n=4 subjects). Representative histogram for β-catenin expression was shown (right). \**P*<0.05 (two-tailed unpaired Student’s *t*-test). **(f, g)** Frequency of IFNγ and IL-10 positive cell number (**f**) and gene expression of *IFNG* and *IL10* by qPCR (**g**). Tregs were stimulated with anti-CD3 and anti-CD28 in the presence of Wnt/β-catenin signaling inhibitor PKF115-584 (PKF), IL-12 (Th1), or IL-12 and PKF115-584 (Th1^+^PKF) (n=4 subjects) \**P*<0.05, \*\**P*<0.01, \*\*\**P*<0.001 (one-way ANOVA with Tukey’s multiple comparisons test). **(h, i)** Relative frequency of IFNγ and IL-10 positive cell number (fold of scramble shRNA/control condition) (**h**) and gene expression of *IFNG* and *IL10* by qPCR (**i**). Tregs were transduced with a non-targeted shRNA or a *CTNNB1* shRNA and cultured in Th0 or Th1 condition for 5 days (**h**; n=7 subjects, **i**; n=5 subjects). \**P*<0.05, \*\**P*<0.01 (one-way ANOVA with Tukey’s multiple comparisons test). Data are representative of two experiments **(e** (middle), and **f)** or are from more than three experiments.

To examine whether the *in vitro* model can recapitulate the *ex vivo* results, we examined active β-catenin levels on each of the Treg subsets after four days of culture with anti-CD3/CD28 stimulation. IFNγ-producing Treg populations (IFNγSP and DP) showed higher active β-catenin expression compared to IL10SP and DN (Fig. 3d), indicating that stabilization of β-catenin is more enhanced in IFNγSP compared to IL10SP under TCR stimulation. IL-12 is an essential cytokine for Th1 differentiation and is known to induce IFNγ-producing pathogenic Tregs under TCR stimulation^8^. We found that upregulation of β-catenin was also observed in IL-12-induced Th1-like Tregs, especially in the IFNγ-producing population (Fig. 3e). To determine if Wnt/β-catenin signaling was necessary for IFNγ production in Th1-like Tregs, we blocked β-catenin signaling with the β-catenin signaling inhibitor, PKF115-584 (PKF). Tregs treated with PKF exhibited a significantly reduced production of IFNγ (Fig. 3f, g). IL-10 production was also suppressed by PKF treatment, albeit less dramatically than IFNγ. To further confirm these results, we knocked down the *CTNNB1* gene in Tregs using short hairpin RNA (shRNA) (Supplementary Fig. 3b) and demonstrated that IL-12-induced IFNγ and IL-10 production was suppressed by silencing of β-catenin (Fig. 3h, i). These data suggest that β-catenin plays a critical role in IFNγ and IL-10 induction in human Tregs, but more profoundly in IFNγ production under TCR stimulation.

### Constitutive activation of β-catenin in Tregs induces *Scurfy-like* autoimmunity

Although the role of Wnt/β-catenin signaling on Tregs has been assessed in different mouse models, the results vary from study to study. To ascertain the physiological relevance of β-catenin signaling in Tregs, we generated Treg-specific β-catenin stabilized mice (*Foxp3^Cre^/β-ctn^ΔEx3^*) by crossing Foxp3-IRES-Cre mice^41^ with *β-ctn^ΔEx3^* mice^42^ (Supplementary Fig. 4a), where the unphosphorylated active form of β-catenin was specifically induced in Tregs. In these *Foxp3^Cre^/β-ctn^ΔEx3^* mice, β-catenin was highly stabilized in Foxp3^+^ Tregs, but not on Foxp3^—^ non-Tregs (Fig. 4a, Supplementary Fig. 4b, c). This mouse model allowed us to assess the role of β-catenin signaling in Foxp3^+^ Tregs more precisely than using Tregs isolated from pan-CD4^+^ T cell specific β-catenin-stabilized mice (*CD4-Cre/β-ctn^ΔEx3^*) mice^30^. Because stabilization of β-catenin in conventional T cells resulted in a highly pro-inflammatory phenotype, there may be a substantial effect from conventional T cells to Tregs in CD4-Cre/*Ctnnb1^ΔEx3^* mice. *Foxp3^Cre^/β-ctn^ΔEx3^* mice spontaneously developed a hunched posture, crusting of the ears, eyelids and tail and showed thymic atrophy, splenomegaly and lymphadenopathy (Fig. 4b). Histologic analysis demonstrated lymphocyte infiltration into several tissues, such as lung, pancreas, liver, and intestine, representing systemic inflammation in *Foxp3^Cre^/β-ctn^ΔEx3^* mice (Fig. 4c). This *scurfy*-like fulminant autoimmunity led to premature death of *Foxp3^Cre^/β-ctn^ΔEx3^* mice within 40 days of birth with 100% penetrance (Fig. 4d).

**Figure 4.**
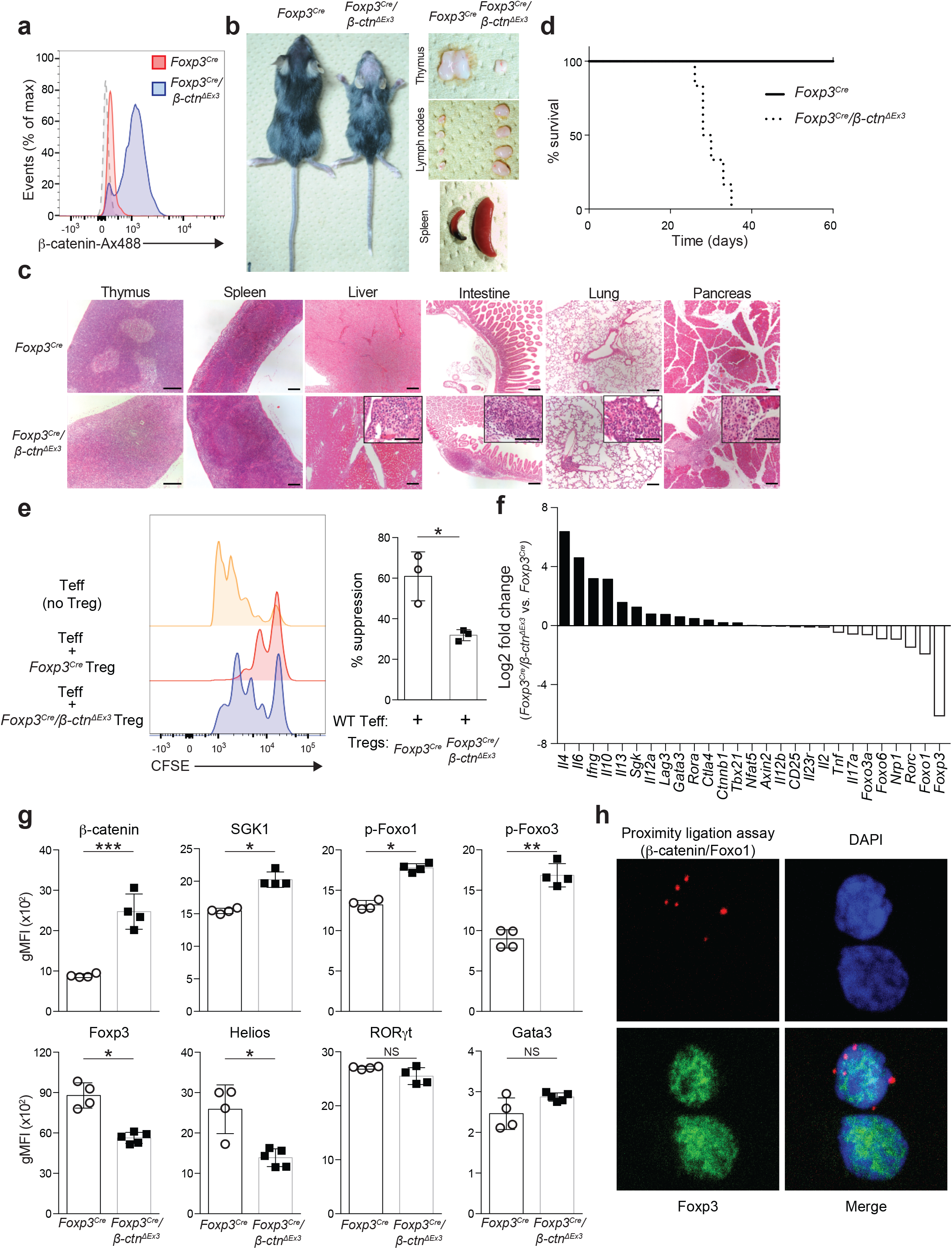
Treg specific activation of β-catenin induces IFNγ secreting dysfunctional phenotype with *Scurfy*-like autoimmunity. **(a)** Flow cytometric analysis of β-catenin in splenic CD4^+^Foxp3^+^ Tregs from Foxp3-IRES-Cre/wild-type mice (*Foxp3^Cre^*) and Foxp3-IRES-Cre/*Ctnnb1^ΔEx3^* (*Foxp3^Cre^/β-ctn^ΔEx3^*) mice. Dashed lane is for isotope control. **(b)** Images of 4-week-old *Foxp3^Cre^* mouse and *Foxp3^Cre^/β-ctn^ΔEx3^* mouse (left). Representative pictures of thymus, peripheral lymph nodes, and spleens isolated from 4 week-old *Foxp3^Cre^* or *Foxp3^Cre^/β-ctn^ΔEx3^* mice. **(c)** Hematoxylin and eosin staining of thymus, spleen, liver, intestine, pancreas, and lung sections from 4 week-old *Foxp3^Cre^* and *Foxp3^Cre^/β-ctn^ΔEx3^* mice. Scale bars, 300 μm in the lower magnification and 150μm in the higher magnification. **(d)** Survival of *Foxp3^Cre^* and *Foxp3^Cre^/β-ctn^ΔEx3^* mice. **(e)** Representative histogram of CFSE dilution for Treg suppression assay. Yellow; *Foxp3^Cre^* Teff only, Blue; *Foxp3^Cre^* Tregs and *Foxp3^Cre^* Teff at 1:1 ratio, Red; *Foxp3^Cre^/β-ctn^ΔEx3^* Tregs and *Foxp3^Cre^* Teff at 1:1 ratio (left). Bar graph shows percentage of suppression (right) (n=3) \**P*<0.05 (two-tailed unpaired Student’s *t*-test). **(f)** Gene expression profile of *Foxp3^Cre^* and *Foxp3^Cre^/β-ctn^ΔEx3^* Tregs by microarray analysis. **(g)** Flow cytometric analysis on peripheral lymph node Tregs from *Foxp3^Cre^* and *Foxp3^Cre^/β-ctn^ΔEx3^* mice. Quantification of gMFI for indicated molecules was shown. (*Foxp3^Cre^*; n=4, *Foxp3^Cre^/β-ctn^ΔEx3^*; n=4 or 5). **(h)** Representative immunofluorescence images of human Tregs with PLA signal for β-catenin-Foxo1 interaction (red) and Foxp3 staining (green). Nuclei were stained with DAPI (blue).

The balance between Tregs and effector T cells is critical to maintain T cell homeostasis both in central and peripheral lymphoid tissue. The percentage of Tregs within thymic CD4^+^ T cells of *Foxp3^Cre^/β-ctn^ΔEx3^* mice remained at the same level as WT mice by the age of 3 weeks and even increased at the age of 5 weeks; however, the number of Tregs in thymus began to decline at the age of 3 weeks in *Foxp3^Cre^/β-ctn^ΔEx3^* mice (Supplementary Fig. 4d). Notably, splenic Tregs showed a significant decrease at an early stage. In contrast, *Foxp3^Cre^/β-ctn^ΔEx3^* mice displayed an increased number of CD4^+^ and CD8^+^ conventional T cells in secondary lymphoid organs (spleen and lymph nodes) and higher expression of effector cytokines such as *IFNG, IL4*, and *IL10*, but not *IL17A* (Supplementary Fig. 4e). Downregulation of *RORC* in both Tregs and conventional CD4^+^ T cells is the opposite for *CD4-Cre/β-ctn^ΔEx3^* mice^30^, highlighting the difference between *CD4-Cre/β-ctn^ΔEx3^* mice and *Foxp3^Cre^/β-ctn^ΔEx3^* mice. To characterize the functional properties of *Foxp3^Cre^/β-ctn^ΔEx3^* Tregs, we examined Helios expression^43^, and found that *Foxp3^Cre^/β-ctn^ΔEx3^* Tregs lost Helios expression compared to *Foxp3^Cre^* Tregs, supporting the unstable and dysfunctional feature of *Foxp3^Cre^/β-ctn^ΔEx3^* Tregs (Supplementary Fig. 4f). These results suggest that forced expression of a stabilized form of β-catenin in Tregs influences their functional stability in the periphery more than in the central compartment.

To examine the suppressive capacity of *Foxp3^Cre^/β-ctn^ΔEx3^* Tregs, we performed an *in vitro* suppression assay. As expected, *Foxp3^Cre^/β-ctn^ΔEx3^* Tregs showed less suppressive activity compared to *Foxp3^Cre^* Tregs (Fig. 4e). Given that the direct interaction of β-catenin with Foxo1 has been reported^44,45^, we noted that the morphological and pathophysiological phenotype of *Foxp3^Cre^/β-ctn^ΔEx3^* mice was similar to that of Treg-specific Foxo1 depletion mice, which exhibited fulminant autoimmunity and disrupted Treg function with aberrant IFNγ expression^46^. To identify transcriptional changes in β-catenin stabilized Tregs, we measured the gene expression signature of Tregs isolated from *Foxp3^Cre^/β-ctn^ΔEx3^* mice by genome-wide DNA microarrays (Fig. 4f). Further assessment with GSEA revealed similar transcriptional profiles between *Foxp3^Cre^/β-ctn^ΔEx3^* Tregs and Foxo1-depleted Tregs (Supplementary Fig. 4h). In agreement with this observation, phosphorylated Foxo1 and Foxo3a were increased in *Foxp3^Cre^/β-ctn^ΔEx3^* Tregs compared to *Foxp3^Cre^* Tregs (Fig. 4g). To determine whether β-catenin and Foxo1 are directly interacting with each other, we performed an *in situ* proximity ligation assay (PLA) on human Tregs and detected the PLA signal in human Tregs (Fig. 4h). Taken together, our results indicate that β-catenin regulates the pro-inflammatory Th1-skewing program in Tregs in concert with the Foxo pathway.

### High salt environment activates the β-catenin/SGK1/Foxo axis and produces IFNγ/IL-10 imbalance

It has been shown previously that the higher sodium concentration detected in tissue interstitium under physiological conditions boosted the induction of Th17 cells and diminished the suppressive capacity of Tregs through activation of SGK1^17, 19^. We recently demonstrated that the PI3K/AKT1/Foxo axis also played a pivotal role in inducing IFNγ-producing dysfunctional Tregs^32^. Furthermore, we observed that p-Foxo1/3a and SGK1 were upregulated in *Foxp3^Cre^/β-ctn^ΔEx3^* Tregs (Fig. 4g). To assess if the SGK1/Foxo axis is activated in human Treg subpopulations, we examined phosphorylated SGK1 (p-SGK1) and phosphorylated Foxo1 (p-Foxo1) levels by flow cytometry and found that both were highly expressed in the IFNγ-producing Treg population *ex vivo* (Supplementary Fig. 5a). Additionally, we demonstrated the direct interaction between β-catenin and Foxo1 in IFNγ-producing human Tregs by using PLA (Supplementary Fig. 5b). These findings prompted us to hypothesize that activation of β-catenin is involved in high salt-induced IFNγ production as an upstream regulator of the SGK1/Foxo axis. Higher expression of active β-catenin, p-SGK1, and p-Foxo1 was observed specifically in the IFNγ-producing human Treg subset under high salt conditions (Fig. 5a), but not in the IL-10-producing subset (Supplementary Fig. 5c). We also confirmed that β-catenin target genes (*AXIN2* and *TCF7*) and TCF-1 protein were upregulated in human Tregs treated with increased salt concentration (Fig. 5b, Supplemental Fig. 5d). Interestingly, additional salt treatment skewed the IL-12 induced, Th1-like Treg to produce more IFNγ and less IL-10, suggesting that the high salt environment might exacerbate the IFNγ-skewing pathogenic Treg signature that resembles the MS-Treg phenotype (Fig. 5c, Supplemental Fig. 5e). To determine if β-catenin activation was necessary for IFNγ induction under high salt conditions, we pharmacologically blocked β-catenin signaling with two different Wnt/β-catenin signaling inhibitors, PKF and IWR-1. PKF or IWR-1 significantly downregulated high salt-induced IFNγ expression in human Tregs (Fig. 5d, Supplementary Fig. 5f). These results were also observed upon genetic knock down of *CTNNB1* by shRNA (Fig. 5e, f). Given that SGK1 is a target of β-catenin signaling^47^, we then tested the impact of inhibiting β-catenin signaling on SGK1 activation. Pharmacological inhibition of β-catenin signaling by PKF or IWR prevented SGK1 phosphorylation under high salt stimulation in human Tregs (Supplementary Fig. 5g). To clarify the role of SGK1/Foxo axis in IFNγ and IL-10 production under high salt conditions, we measured the production of IFNγ and IL-10 and Foxo1 phosphorylation in the presence or absence of SGK1 inhibitor (GSK650394) in human Tregs. GSK650394 treatment ameliorated high salt-induced IFNγ production and Foxo1 phosphorylation, but had no impact on IL-10 production (Supplementary Fig. 5h, i). These results suggest that β-catenin positively regulates salt-induced IFNγ expression through activation of the SGK1/Foxo axis. We also extended the analysis to human effector T cells (Teff, CD4^+^ CD25^−^) and Jurkat T cells. Both of these displayed active β-catenin signaling under high salt condition (Supplementary Fig. 6a). In Teff cells there was also an imbalance of IFNγ and IL-10 production in agreement with our Treg data (Supplementary Fig. 6b). In addition, to further define the action of β-catenin/SGK1/Foxo1 axis in high salt conditions, we generated β-catenin-depleted human Jurkat T cells by using CRISPR/Cas9 technology and examined whether SGK1 and Foxo1 phosphorylation were regulated by β-catenin during high salt exposure. High salt-induced SGK1 and Foxo1 phosphorylation were attenuated in β-catenin knockout Jurkat T cells (Supplementary Fig. 6c, d). These data, along with the evidence from non-immune cells^48^, supports our hypothesis that the β-catenin/SGK1/Foxo1 axis is activated by high salt stimulation.

**Figure 5.**
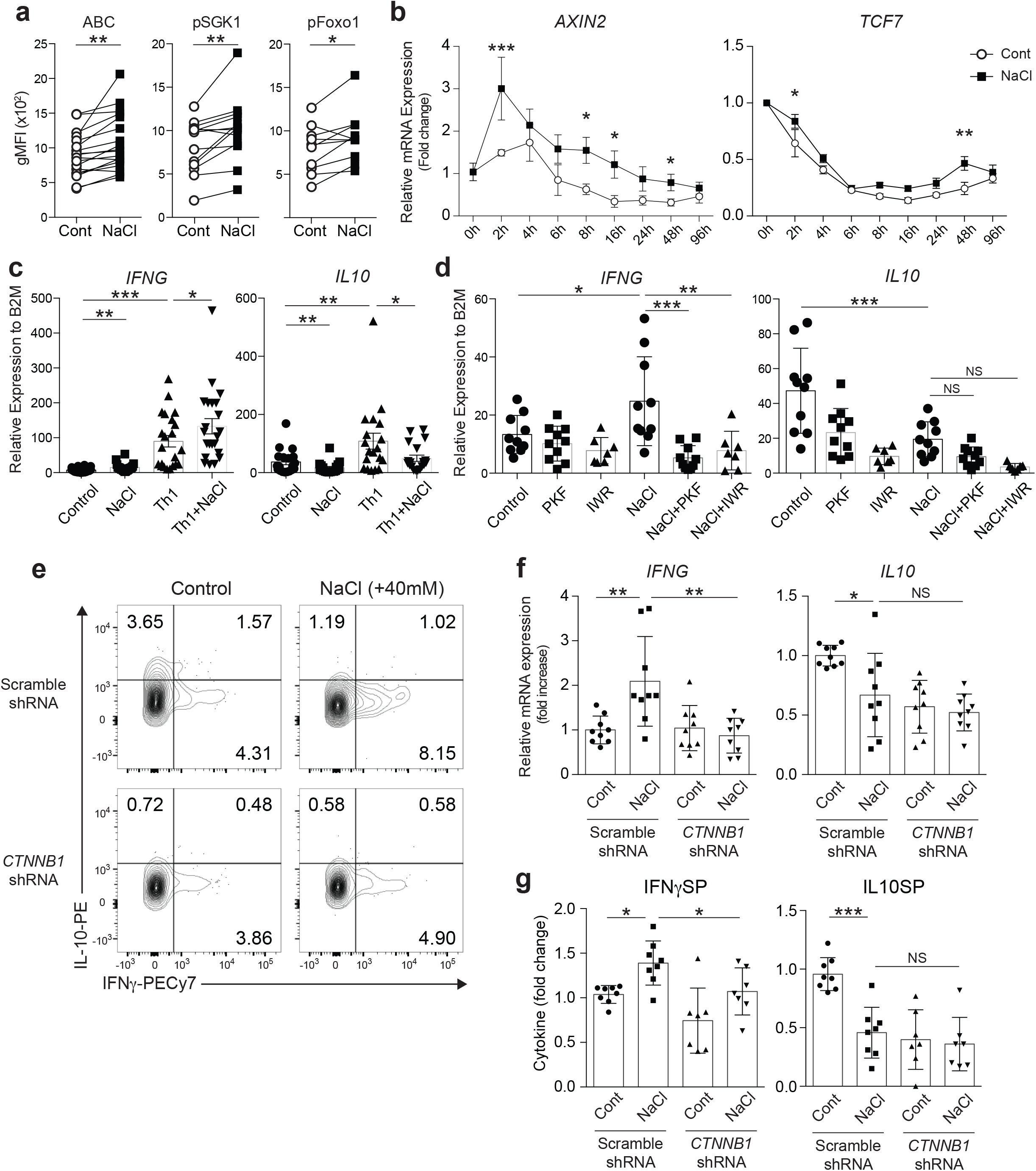
High salt environment induces β-catenin signal activation and IFNγ/IL-10 cytokine imbalance. **(a)** Flow cytometric analysis of ABC, phospho-SGK1 (Thr256), and phospho-Foxo1 (Ser256) expression in human IFNγ-producing Tregs. Tregs were stimulated with anti-CD3 and anti-CD28 in the presence (NaCl) or absence (Control) of additional 40 mM NaCl for 96 h followed by 4 h PMA/iomomycin stimulation (n=13-18 subjects). \**P*<0.05, \*\**P*<0.01 (two-tailed unpaired Student’s *t*-test). **(b)** mRNA expression kinetics for Wnt/β-catenin target genes (*AXIN2* and *TCF7*) from 9 time points were plotted and each dots represent the average of 4 different experiments. \**P*<0.05, \*\**P*<0.01, \*\*\**P*<0.001 (two-way ANOVA with Sidak’s multiple comparisons test). **(c)** *IFNG* mRNA expression in human Tregs cultured in Th0 or Th1 condition in the presence (NaCl) or absence (Control) of additional 40 mM NaCl for 96 h (n=19 subjects). \**P*<0.05, \*\**P*<0.01, \*\*\**P*<0.001 (one-way ANOVA with Tukey’s multiple comparisons test). **(d)** *IFNG* mRNA expression in human Tregs stimulated in the presence (NaCl) or absence (Control) of additional 40 mM NaCl with and without Wnt/inhibitor PKF115-584 (PKF) or IWR-1 (IWR) for 96 h (n=7-10 subjects). \**P*<0.05 (one-way ANOVA with Tukey’s multiple comparisons test). **(e)** Representative flow cytometric analysis of IFNγ and IL-10 production in human Tregs transduced with a non-targeted shRNA or a *CTNNB1* shRNA and cultured in the normal media (Control) or media supplemented with additional 40 mM NaCl (NaCl) for 96 h. **(f)** *IFNG* and *IL10* mRNA expression on Tregs, and **(g)** frequency of IFNγ and IL-10 producing Tregs relative to control/scramble shRNA condition were shown. Tregs were treated as in **(e)** (**f**; n=9 subjects, **g**; n=8 subjects). \**P*<0.05, \*\**P*<0.01, \*\*\**P*<0.001 (one-way ANOVA with Tukey’s multiple comparisons test).

We next explored the molecular mechanisms underlying high salt-induced β-catenin activation. First, we examined whether Wnt ligands play a role in this aberrant activation of β-catenin signaling in a high salt environment. We used Frizzled-8 FC chimera protein (Fzd8-FC), which is known to act as a scavenger of Wnt ligands, to inhibit the effect of Wnt ligands on Tregs. Activation of β-catenin assessed by ABC level or production of IFNγ and IL-10 were not affected by Fzd8-FC treatment in control or high salt conditions, suggesting that the role of Wnt ligands in high salt-induced activation of β-catenin signaling was dispensable (Supplementary Fig. 7a, b). Although a salinity stress sensor has not been fully described in mammalian cells, a number of pathways contributing to the salt stress response have been identified^48, 49^. Within these pathways, we focused on AKT kinase because it is well known to regulate β-catenin signaling via direct phosphorylation of β-catenin^50^ or indirectly through GSK3β, which is a negative regulator of β-catenin^51^. Moreover, AKT is known to play a prominent role on Th1 differentiation, and we recently reported that AKT1 acted as a potent inducer for IFNγ-producing, Th1-like Tregs and that AKT1 inhibition could recover the dysfunctional MS-Treg phenotype^32^. Indeed, the PI3K-AKT pathway was highly enriched in the IFNγ-producing Treg subset and AKT phosphorylation at S473 was increased in IFNγ-producing Tregs (Supplementary Fig. 7c, d). These results prompted us to hypothesize that activation of AKT kinase regulates β-catenin signaling during chronic high salt stress conditions. To examine the effect of chronic high salt stimulation independently of TCR signaling, we took advantage of the Jurkat T cell line, that can be maintained without TCR stimulation. We then investigated whether β-catenin could be directly activated by AKT by examining AKT-specific phosphorylation of β-catenin (Ser522), which stabilizes β-catenin^50^. Phosphorylation of β-catenin (Ser522) was increased in a high salt environment and this effect was reversed by the AKT inhibitor MK2206, indicating that activation of AKT is responsible for stabilizing β-catenin during high salt stimulation^48^ (Supplementary Fig. 7e). Furthermore, we demonstrate that phosphorylation of GSK3β at Ser9, which is an important site of phosphorylation by AKT^52^, was increased by high salt stimulation (Supplementary Fig. 7f). We also confirmed that both p-AKT and p-GSK3β levels were not affected by silencing β-catenin (Supplemental Fig. 7f), suggesting that both of them act upstream of β-catenin. These data indicate that AKT regulates β-catenin activation in both direct and indirect mechanisms under high salt conditions.

### A high salt-induced PTGER2-β-catenin loop leads to imbalance between IFNγ and IL-10

Both IFNγ and IL-10 are upregulated in IL-12-induced Th1-like Tregs in a β-catenin-dependent manner (Fig. 3f-i). However, IL-10 expression was significantly suppressed by high salt treatment, contrary to IFNγ. In fact, the β-catenin/SGK1/Foxo axis was not activated in IL-10SP after high salt treatment (Supplementary Fig. 5c). Additionally, the effect of high salt on IL-10 production could not be explained by activated β-catenin signaling (Fig. 5e, f, Supplementary Fig. 5f). Thus, we hypothesized that there might exist a factor that can be uniquely induced in the high salt environment but not in IL-12-driven Th1 conditions, resulting in IL-10 inhibition. This was addressed by comparing gene expression profiles of Tregs between control media and IL-12 supplemented media (Th1), and also between control media and NaCl supplemented media (NaCl). Among the group of differentially expressed genes in each comparison, we identified six genes that were upregulated in high salt conditions but downregulated in Th1 conditions, and four genes that were regulated in the opposite direction, which are potentially able to account for the high salt-induced IFNγ/IL10 imbalance (Fig. 6a).

**Figure 6.**
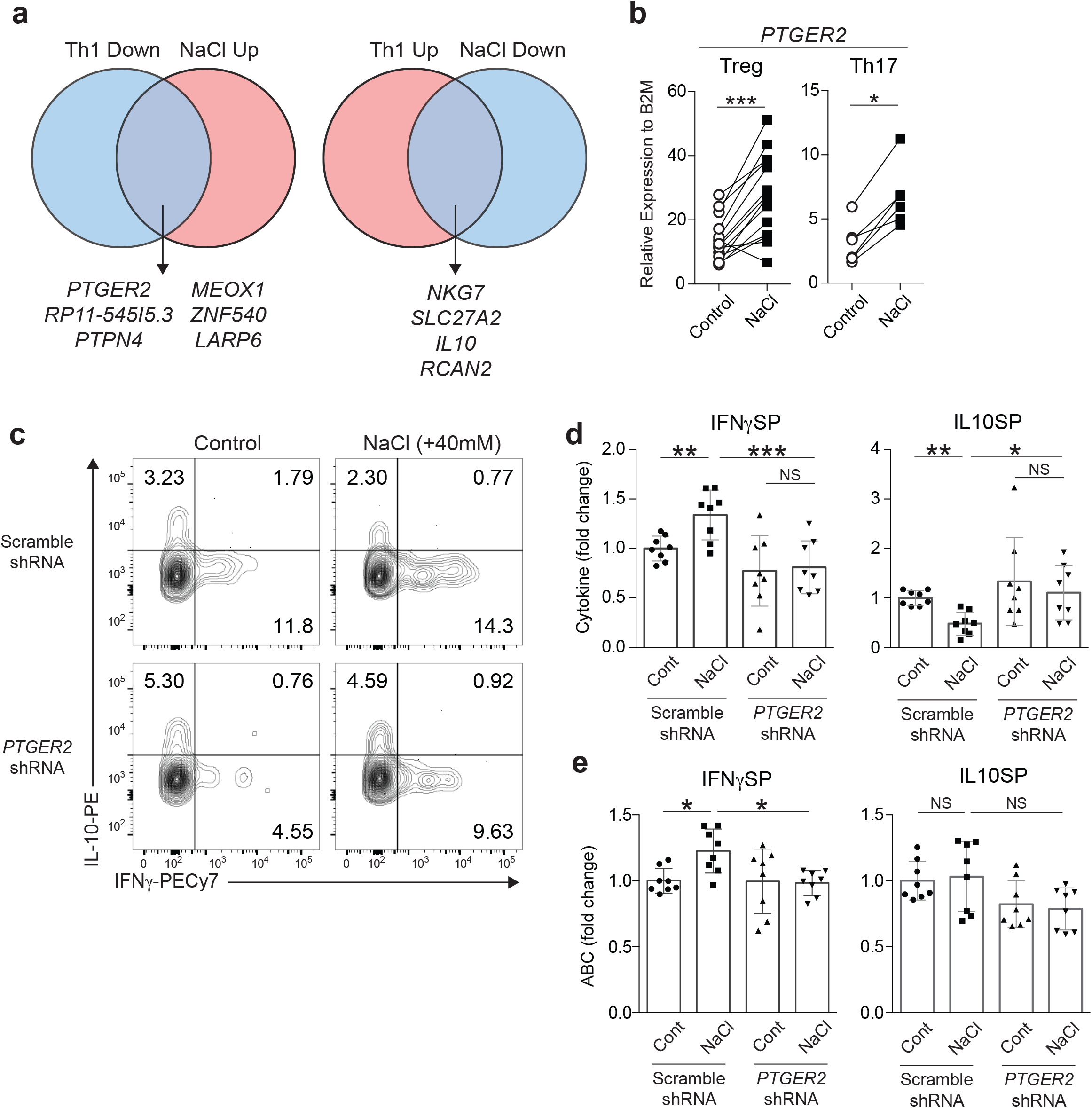
PTGER2 is a unique factor regulating IFNγ and IL-10 in conjunction with β-catenin under high salt condition. **(a)** Venn diagrams showing the overlapped genes between the genes upregulated in NaCl treatment (NaCl Up) and downregulated in Th1 condition (Th1 Down) (left), and between the genes downregulated in NaCl treatment (NaCl Down) and upregulated in Th1 condition (Th1 Up) (right). **(b)** *PTGER2* mRNA expression in human Tregs (left) and Th17 cells (right). Tregs were stimulated with anti-CD3 and anti-CD28 in the normal media (Control) or media supplemented with additional 40 mM NaCl (NaCl) for 96 h (n=14 subjects). Naïve CD4^+^ T cells were cultured in the normal Th17 condition (Control) or Th17 condition supplemented with additional 40 mM NaCl (NaCl) for 72 h (n=6 subjects). \**P*<0.05, \*\*\**P*<0.001 (two-tailed Student’s *t*-test). **(c)** Representative flow cytometric analysis of IFNγ and IL-10 production in human Tregs transduced with a scramble shRNA or a *PTGER2* shRNA and cultured in the normal media (Control) or media supplemented with additional 40 mM NaCl (NaCl) for 96 h. **(d)** Relative frequency of IFNγ and IL-10 producing Tregs, and **(e)** relative expression level of ABC in Tregs were shown. Tregs were treated as in **(c)** (**d, e**; n=8 subjects). \**P*<0.05, \*\**P*<0.01, \*\*\**P*<0.001 (one-way ANOVA with Tukey’s multiple comparisons test).

Prostaglandin E receptor 2 (PTGER2) is in the family of G protein-coupled receptors, and acts as one of the receptors for prostaglandin E2 (PGE2). PGE2-PTGER2 signaling increases intracellular cAMP, which in turn activates PKA signaling, dampening T cell activation^53^. However, the signal through PTGER2 is also known to regulate the production of cytokines in a context-dependent manner^54–58^, and the strength of PI3K-AKT signaling has been reported as an important component affecting the action of PTGER2 on cytokine production, especially IFNγ production in T cells^59^. Since we have observed a role for PTGER2 in promoting the pathogenic phenotype by modulating IFNγ/IL10 balance in Th17 cells^56^ and high salt treatment induces a pathogenic Th17 cell signature, we hypothesized that PTGER2 regulates the IFNγ/IL-10 balance in salt stimulated Tregs. Indeed, *PTGER2* was upregulated after high salt treatment in human Tregs and Th17 cells (Fig. 6b) and was highly expressed in IFNγSP compared to IL10SP (Supplementary Fig. 8a).

Given the evidence of a positive relationship between β-catenin signaling and PTGER2^60, 61^, we investigated whether β-catenin and PTGER2 build an autoregulatory loop during chronic high salt exposure. To understand the link between β-catenin and PTGER2 in the high salt environment, we used Jurkat T cells to demonstrate that high salt-induced PTGER2 was suppressed by genetic deletion of *CTNNB1* (Supplementary Fig. 8b). We also confirmed that PTGER2 knockdown could partially ameliorate high salt-induced β-catenin activation (Supplementary Fig. 8c). These results suggest the presence of a β-catenin-PTGER2 feed-forward loop under high salt conditions. We then examined whether the PTGER2-β-catenin loop contributes to the IFNγ-IL-10 balance in human Tregs under high salt conditions. To this end, human Tregs were transduced with shRNA directed against PTGER2 and stimulated in the presence and absence of additional NaCl. PTGER2 silencing abolished the high salt-induced imbalance of IFNγ and IL-10 production (Fig. 6c, d). Moreover, PTGER2 knockdown eliminated the high salt-induced increase of ABC in IFNγSP, while it did not affect ABC level in IL10SP, suggesting that induction of IFNγ by high salt conditions depended on the PTGER2-β-catenin loop, but IL-10 suppression by high salt is dependent on PTGER2 *per se* but not on β-catenin.

PTGER2 is known to regulate cytokine production in a context-dependent manner^58^, and the strength of PI3K-AKT signaling has been reported as one of the important components affecting the action of PTGER2 on cytokine production, especially in IFNγ production on conventional CD4^+^ T cells^59^. Based on this evidence, we hypothesized that the high salt-induced PTGER2-β-catenin loop could be amplified in cells where AKT is activated, such as in IFNγ producing Tregs (IFNγSP), but not in the cells with lower AKT activity, such as IL-10 producing Tregs (IL10SP). We then tested the impact of modulating AKT signaling on IFNγ and IL-10 production in Tregs under high salt conditions via increasing CD28 co-stimulation. High salt-induced IFNγ production was boosted by strengthening AKT signaling with higher CD28 co-stimulation (Supplementary Fig. 8d)^59^. By contrast, high salt-induced IL-10 inhibition was not altered. Together, these data indicated that high salt induces a positive feedback loop between β-catenin and PTGER2 in conjunction with activated AKT status, resulting in amplification of IFNγ production in Tregs.

### Stabilized β-catenin is observed in Tregs from mice fed a high salt diet and MS patients

To examine if β-catenin is stabilized under high salt conditions *in vivo*, we fed wild type mice with either a normal salt diet (NSD), containing 0.4% of NaCl, or a high salt diet (HSD), containing 4% NaCl, and assayed β-catenin expression on Tregs. We found that β-catenin and phosphorylated Foxo1/3a/4 were increased in Tregs from high salt diet mice (Fig. 7a). Next, we determined if β-catenin is more stabilized in MS-Tregs as compared to Tregs from healthy subjects. The level of active β-catenin in IFNγ producing Tregs was increased in MS patients compared to healthy subjects (Fig. 7b). We also found a positive correlation between IFNγ production and the level of active β-catenin in MS-Tregs but not in healthy subjects (Fig. 7c). Furthermore, to investigate the link between *PTGER2* expression and active β-catenin level or *IFNG* expression in MS-Tregs, we assessed the expression of these factors in Tregs from healthy subjects and MS patients. Notably, higher expression of *PTGER2* and active β-catenin level correlated with *IFNG* expression in MS-Tregs but not in Tregs from healthy subjects (Fig. 7d, e). These findings provide *in vivo* evidence to support our hypothesis that the PTGER2-β-catenin loop plays an important role in the salt-induced malfunction of Tregs and links this salt signature to the pathogenic profile of MS-Tregs.

**Figure 7.**
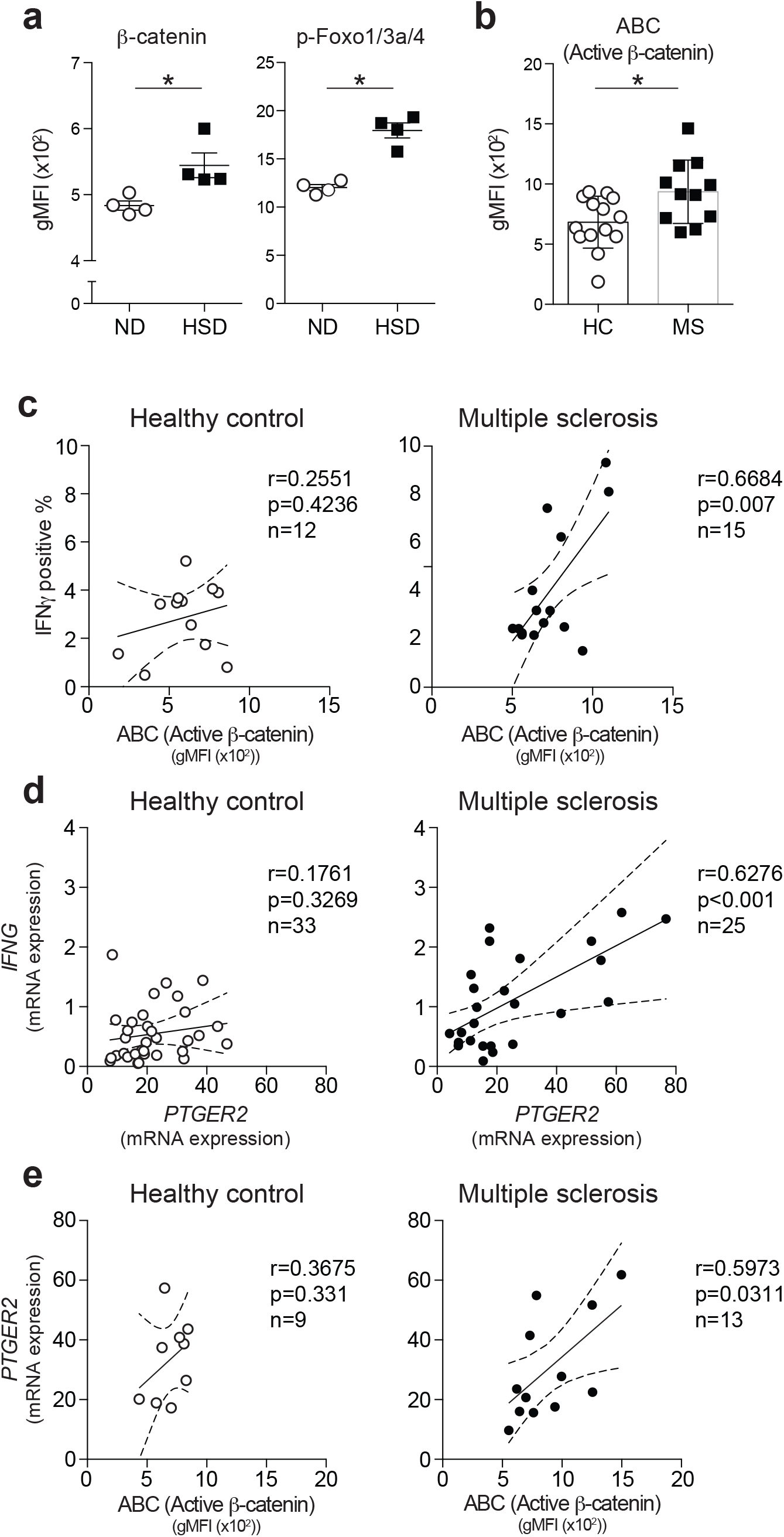
Stabilized β-catenin associated with IFNγ and PTGER2 expression in Tregs from MS patients. **(a)** Flow cytometric analysis of Tregs from the mesenteric lymph nodes of wild type mice fed a normal diet (ND) or a high-salt diet (HSD) for 3 weeks. Quantification of gMFI for β-catenin and p-Foxo1/3a/4 were shown (ND; n=4, HSD; n=4). \**P*<0.05 (two-tailed unpaired Student’s *t*-test). **(b)** Flow cytometric analysis of ABC level in *ex vivo* Tregs of healthy controls and MS patients (HC; n=14 subjects, MS; n=11 subjects). Correlation plots **(c)**; between the percentage of IFNγ-producing Tregs and gMFI of Active β-catenin (ABC), **(d)**; between *IFNG* and *PTGER2* mRNA expression, **(e)**; between ABC level and *PTGER2* mRNA expression level in healthy subjects and MS patients. Linear regression is shown with 95% confidence interval (dotted lines). Correlation statistics by two-tailed Spearman rank correlation test.

## Discussion

Loss of Foxp3^+^ regulatory T cell function is associated with a number of autoimmune diseases including MS^8^ and type 1 diabetes^9^, and clinical trials involving adoptive transfer of autologous Tregs are now in progress. We and others have shown that IFNγ expression correlates with loss of Treg function, and that blocking IFNγ can partially restore *in vitro* suppressor function. Moreover, a high salt environment skews Tregs into an IFNγ-producing dysfunctional state. Here, we present data demonstrating that not only induction of IFNγ, but also reduction of IL-10, is observed in Tregs from MS patients as a shared signature with high salt conditions. Using RNA-seq based transcriptional profiling of IFNγ and/or IL-10 producing human Tregs, we identified β-catenin as an upstream regulator for IFNγ and IL-10 production. Treg-specific activation of β-catenin resulted in scurfy-like lethal autoimmunity accompanied by Treg dysfunction and excessive IFNγ expression. Furthermore, β-catenin is stabilized under high salt conditions and acts upstream of the SGK1/Foxo axis, accounting for the pathogenic phenotype. Our results demonstrate a novel role of β-catenin as a regulatory molecule for Treg functional plasticity and also provide molecular mechanisms that link an environmental factor to autoimmune disease (Supplementary Fig. 9).

It has become apparent that Tregs display heterogeneity and plasticity in function and cytokine profile and an IFNγ-producing phenotype has been observed in autoimmune diseases, such as MS^8^ and T1D^9^. These Th1-like Tregs are dysfunctional and are considered to be important in disease pathophysiology. In contrast, IL-10 producing Tregs appear to play a protective role in autoimmunity, given that IL-10 is an anti-inflammatory cytokine and defects in IL-10 are known to induce autoimmunity^11^. However, the molecular mechanisms that control IL-10 production in Tregs and the mechanisms that regulate the balance between IFNγ and IL-10 are poorly understood. Moreover, with clinical trials of autologous Treg infusions in patients with autoimmune disease, identification of targetable pathways to alter the dysfunctional Treg state may be of clinical utility.

In our experiments, transcriptional profiling of human Treg subsets based on IFNγ and IL-10 production provided new insights into Treg heterogeneity. We identify β-catenin as a key regulator identified by upstream regulator analysis and demonstrate its role in skewing Tregs into an IFNγ-producing dysfunctional state in human Tregs and in murine models. Of note, we also identified additional upstream regulators, either previously reported or of interest for future studies: *MYB, SATB1, NFATC2, KLF2, ZBTB16*, and *DDIT3* were all found to be significant upstream regulators across the Treg subpopulations. Treg-specific roles for *MYB*^33^, *SATB1*^34^, *NFATC2*^35^, *and KLF2*^36^ have already been demonstrated in maintaining Treg functional properties, supporting the validity of our upstream regulator analysis. Given that these factors have a capacity to interact with β-catenin, for example, β-catenin can form a complex with SATB1 and modulate Th2 cell differentiation^62^, it is likely that these factors can interact to establish a complex regulatory system in Treg homeostasis.

Although several studies have demonstrated the role of β-catenin in Tregs, it is still unknown how β-catenin contributes to Treg function and the effector cytokine signature. One study demonstrated β-catenin as an anti-inflammatory factor in the context of generating long-lived suppressive Tregs via anti-apoptotic gene induction^28^, and two previous studies showed that activation of β-catenin provokes Treg dysfunction, leading to exaggerated colitis in a murine model^29,30^. We show that Treg-specific stabilization of β-catenin resulted in loss of suppressive properties of Tregs and induced a lethal *scurfy*-like phenotype in mice. Tregs isolated from *Foxp3^Cre^/β-ctn^ΔEx3^* mice had significant upregulation of effector cytokines, such as IFNγ, IL-4, IL-10, but not IL-17A. This result is consistent with a model, supported by our transcriptional analysis on human Treg subsets, where β-catenin plays a role in IL-10 as well as IFNγ production. While a Th17-like signature was previously reported as a result of activation of β-catenin in Tregs, the examination of cytokine signature and functional experiments were all performed on Tregs isolated from *CD4-Cre/Ctnnb1^ΔEx3^* mice^30^. Given that conventional CD4^+^ T cells in *CD4-Cre/Ctnnb1^ΔEx3^* mice displayed a robust pro-inflammatory phenotype, the properties of Tregs in those mice can be affected by their altered cytokine microenvironment, and it is not surprising that our results with Treg-specific genetic intervention using *Foxp3-Cre* mice provide different results as compared to *CD4-Cre*-based intervention. Our *Foxp3-Cre*-based Treg specific intervention provides direct evidence for the role of β-catenin in Tregs.

The incidence of autoimmune diseases has been increasing in the last half century, which cannot be explained by genetic adaptation. Thus, there is great interest in studying the interplay between genetic risks and environmental triggers^63^. Among several environmental triggers, a high salt diet may be one factor contributing to the increasing incidence of autoimmune diseases, though this as yet requires further epidemiologic investigations^64^. There is growing evidence that the excess uptake of salt can modulate both the innate and adaptive immune system^65^. We previously observed that higher salt concentration has direct effects on the induction of Th17 cells and the stability of Tregs, manifest by aberrant IFNγ production^19^. Of note, we also found that high salt-exposed Tregs lose IL-10 expression. This IFNγ/IL-10 imbalance was observed in MS-Tregs, suggesting that the salt-induced Treg signature may overlap with the MS pathogenic profile. The importance of IFNγ/IL-10 balance in the context of salt-induced immune alteration was supported by a previous study showing that increased sodium content in colon tissue of HSD mice resulted in excessive inflammation in IBD models^21^. Interestingly, β-catenin signaling was activated not only in Tregs but also in Th17 cells under stimulation with high salt (data not shown). Furthermore, here we demonstrate that PTGER2 accounts for high salt-induced IFNγ/IL-10 imbalance in Tregs by creating a positive feed forward loop with β-catenin. Notably, previous studies have clarified the interaction between PGE2, a ligand for PTGER2, and Wnt/β-catenin signaling in colorectal carcinogenesis^60, 61, 66^. It has also been reported that Tregs can produce PGE2^67^ and that PGE2 is enriched in EAE lesions^68^. Therefore, PGE2-PTGER2 signaling could be amplified in Tregs under high salt conditions and also in the MS lesion. However, the role of PGE2 in EAE and MS remains unclear and further investigation is needed. Given that emerging studies have highlighted the role of PGE2 signaling and Wnt/β-catenin signaling in mucosal tolerance, our finding of a high salt-induced PTGER2/β-catenin axis may provide new insights into the regulation of gut immune homeostasis. Since β-catenin activation is reported to induce tolerogenic dendritic cells and PTGER2 has a context-dependent role on innate cells, future studies with more detailed characterization of PTGER2-β-catenin signaling dynamics in mucosal immune homeostasis are warranted.

In summary, we provide genome-wide transcriptomic profiles of human *ex vivo* Treg subpopulations, which unveil the heterogeneity of Tregs in terms of IFNγ and IL-10 production. Further analysis and biological validation identify β-catenin as a key molecule in maintaining Treg function and production of IFNγ/IL-10 cytokines in both human and murine Tregs. Moreover, we uncover the unexpected role of PTGER2 in driving the imbalance of IFNγ/IL-10 concomitant with activation of β-catenin signaling under high salt conditions. Lastly, we show that Tregs from MS patients display positive correlations among IFNγ production, *PTGER2* expression, and active β-catenin level, which are not observed in Tregs from healthy subjects (Fig. 7c-e). Thus, our study in humans with autoimmune disease and confirmed in mouse models indicates that the PTGER2-β-catenin axis serves as a bridge between environmental factors and autoimmune disease by modulating Treg properties.

## Methods

### Study subjects

Peripheral blood was drawn from healthy individuals and patients with MS after informed consent and approval by the Institutional Review Board at Yale University. The patients were diagnosed with either Clinically Isolated Syndrome (CIS) or Relapsing-Remitting MS (RRMS) by 2010 MacDonald Criteria and were not treated with any immunomodulatory therapy at the time of the blood draw. All experiments conformed to the principles set out in the WMA Declaration of Helsinki and the Department of Health and Human Services Belmont Report. Clinical characteristics of evaluated MS patients are listed in Supplementary Table 1.

### Mice

C57BL/6J mice were purchased from the Jackson Laboratory or CLEA Japan. FIC mice^41^, and *Ctnnb1*^ΔEx3^ mice^42^ have been described, and mice backcrossed into the C57BL/6J strain were used. *Foxp3^Cre^/β-ctn^ΔEx3^* mice were studied at 3-5 weeks of age. For high salt diet (HSD) experiments, six-week-old male wild type mice were fed with normal chow (control group) or sodium-rich chow containing 4% NaCl (Research Diets; HSD group) with normal tap water for 3 weeks. Cells were isolated from the spleen and/or mesenteric lymph nodes and Foxp3^+^Tregs were analyzed by flow cytometry. All experiments were approved by the University of Tokyo Ethics Committee for Animal Experiments and strictly adhered to the guidelines for animal experiments of the University of Tokyo.

### Human T cell isolation and culture

Peripheral blood mononuclear cells (PBMCs) were isolated from donors by Ficoll-Paque PLUS (GE Healthcare) or Lymphoprep (Stemcell) gradient centrifugation. Total CD4^+^ T cells were isolated by negative magnetic selection using a CD4 T cell isolation kit (Stemcell) and CD4^+^CD25^hi^CD127^low-neg^CD45RO^+^ Tregs were sorted on a FACS Aria (BD Biosciences). Tregs were cultured in RPMI 1640 medium supplemented with 5% Human serum, 2 nM L-glutamine, 5 mM HEPES, and 100 U/ml penicillin, 100 μg/ml streptomycin, 0.5 mM sodium pyruvate, 0.05 mM nonessential amino acids, and 5% human AB serum (Gemini Bio-Products). 96-well round bottom plates (Corning) were pre-coated with anti-human CD3 (UCHT1) (1-2 μg/ml) and used for Treg *in vitro* culture with soluble anti-human CD28 (28.2) (1-5 μg/ml) (BD Bioscience) and human IL-2 (50 U/ml). Human IL-2 was obtained through the AIDS Research and Reference Reagent Program, Division of AIDS, National Institute of Allergy and Infectious Diseases (NIAID), National Institutes of Health (NIH). Th1-Tregs induced with human recombinant IL-12 (20 ng/ml) (R&D). The Wnt/β-catenin inhibitor PKF115-584 (Tocris) was used at 200 nM and IWR-1 (Tocris) was used at 20 μM. SGK1 inhibitor GSK650394 (Tocris) was used at 10 μM. AKT inhibitor MK2206 (Tocris) was used 5 μM. Frizzled 8 FC chimera protein (R&D) was used at 500 ng/ml.

### Suppression assay

CD4^+^CD25^+^ Tregs and CD4^+^CD25^−^ T responder cells were isolated from spleen and lymph node from *Foxp3^Cre^* mice or *Foxp3^Cre^/β-ctn^ΔEx3^* mice by using CD4^+^CD25^+^ Regulatory T Cell Isolation Kit (Miltenyi Biotec). T responder cells were labeled with CFSE and then co-cultured with Tregs (5 x 10^4^) at 1:1 ratio in RPMI 1640 medium supplemented with 10% FBS (HyClone), 50 μM 2-Mercaptoethanol (Sigma-Aldrich), 1x GlutaMAX, 50□U/ml penicillin, and 100□μg/ml streptomycin with Dynabeads Mouse T-Activator CD3/CD28 at 2:1 bead-to-cell ratio. The proliferation of T responder cells was determined at day 4 by FACS on a Verse instrument (BD Bioscience).

### Quantitative PCR

Total RNA was extracted using RNeasy Micro Kit (QIAGEN), or ZR-96 Quick-RNA kit (Zymo Research), according to the manufacturer’s instructions. RNA was treated with DNase and reverse transcribed using TaqMan Reverse Transcription Reagents (Applied Biosystems) or SuperScript IV VILO Master Mix (Invitrogen). cDNAs were amplified with Taqman probes (Taqman Gene Expression Arrays) and TaqMan Fast Advanced Master Mix on a StepOne Real-Time PCR System (Applied Biosystems) according to the manufacturer’s instructions. mRNA expression was measured relative to *B2M* expression.

### Flow cytometry analysis

Single-cell suspensions were prepared from PBMCs or mouse tissues and stained with fixable viability dye for 10 min at RT, followed by staining with surface antibodies for 30 min at 4°C. For intracellular staining, cells were fixed and permeabilized with the Foxp3 Fix/Perm buffer set (eBioscience) for 1 h at 4°C, followed by staining with intracellular antibodies. For cytokine staining, cells were stimulated with PMA (50 ng/ml) and ionomycin (for *ex vivo* Tregs; 1000 ng/ml, for *in vitro* cultured Tregs; 250 ng/ml) in the presence of GolgiPlug (BD Bioscience) for 4 h at 37°C. Antibodies and reagents used for flow cytometric analysis are listed in Supplementary Table 2. Stained samples were analyzed with a BD FACS Verse or an LSR Fortessa flow cytometer (BD Bioscience). Data were analyzed with FlowJo software (Treestar).

### RNA-seq library preparation and data analysis

#### Preparation of cells for RNA-seq

For the *ex vivo* Treg subpopulations, CD4^+^CD25^hi^CD127^neg-low^CD45RO^+^ memory Tregs from healthy donors were sorted and immediately stimulated with PMA (50 ng/ml) and iomomycin (1000 ng/ml) for 4 h. By combining IFNγ secretion assay (APC) and IL-10 secretion assay (PE) (Miltenyi), Tregs were labeled based on the expression of IFNγ and IL-10. To avoid RNA degradation, cells were kept in CellCover (Anacyte) before a second round of sorting. For *in vitro* cultured Tregs in Th1 or high salt conditions, mTregs were cultured in each condition for four days as described. Cells were harvested and immediately processed for cDNA preparation. Samples were collected from four healthy subjects for identification of the Th1 signature and five healthy subjects for identification of the high salt signature.

#### cDNA and library preparation and sequencing

cDNAs were generated directly from resorted and harvested cells using the SMART-Seq v4 Ultra Low Input RNA Kit for sequencing (Takara/Clontech). Barcoded libraries were generated by the Nextera XT DNA Library Preparation kit (Illumina) and sequenced with a 2 x 100 bp paired-end protocol on the HiSeq 2000 Sequencing System (Illumina).

#### RNA-seq data analysis

RNA-seq analysis was performed using Partek flow (v6.6). First, RNA-seq reads were trimmed and mapped to the hg19 genome reference using STAR (2.5.0e). Aligned reads were quantified to the gene level using Partek’s E/M algorithm and gene annotation from Ensembl release 75. Gene-level quantitations were normalized by dividing the gene counts by the total number of reads, following by addition of a small offset (0.001). The offset was added to enable log2 transformation and the value of the offset was determined by exploring the data distribution. Differential expression was assessed by fitting Partek’s log-normal model with shrinkage (comparable in performance to limma-trend). Genes having geometric mean below 1.0 were removed from the analysis.

For *ex vivo* Treg subpopulation data, differentially expressed genes (Fold change > 1.5, *P* value < 0.05) were used for functional analysis using IPA and upstream regulator analysis (https://www.qiagenbioinformatics.com/products/ingenuity-pathway-analysis/). Gene set enrichment analysis was performed on normalized gene expression counts of RNA-seq data or microarray data as described previously.

For *in vitro* Th1-Treg and high salt Treg data, differentially expressed genes (Fold change > 2, *P* value < 0.05) were used.

### Microarray

For the Oligo DNA microarray analysis, total RNA samples were extracted from sorted CD4^+^CD25^hi^Tregs of *Foxp3^Cre^* mice or *Foxp3^Cre^/β-ctn^ΔEx3^* mice. Microarray analysis was performed with a 3D-Gene Mouse Oligo chip 24k (Toray Industries Inc., Tokyo, Japan). Total RNA was labeled with Cy5 by using the Amino Allyl MessageAMP II aRNA Amplification Kit (Applied Biosystems). The Cy5-labeled aRNA pools were mixed with hybridization buffer, and hybridized for 16 h. The hybridization signals were obtained by using a 3D-Gene Scanner and processed by 3D-Gene Extraction software (Toray Industries Inc.). Detected signals for each gene were normalized with the global normalization method (the median of the detected signal intensity was adjusted to 25).

### Histology

Mouse tissues were fixed in Ufix (Sakura Finetek Japan) and embedded in paraffin. 6-μm tissue sections were stained with haematoxylin and eosin.

### Lentiviral transduction for shRNA gene silencing and CRISPR/Cas9-mediated gene deletion

Lentiviral plasmids encoding shRNAs were obtained from Sigma-Aldrich and all-in-one vectors carrying *CTNNB1* sgRNA/Cas9 with GFP reporter were obtained from Applied Biological Materials. Each plasmid was transformed into One Shot^®^ Stbl3™ chemically competent cells (Invitrogen) and purified by ZymoPURE plasmid Maxiprep kit (Zymo research). Lentiviral pseudoparticles were obtained after plasmid transfection of 293FT cells using Lipofectamine 2000 (Invitrogen). The lentivirus-containing media was harvested 48 or 72 h after transfection and concentrated 40 - 50 times using Lenti-X concentrator (Takara Clontech). Sorted Tregs were stimulated with plate-bound anti-CD3 (1 μg/ml) and soluble anti-CD28 (1 μg/ml) for 24 h and transduced with lentiviral particles by spinfection (1000 x *g* for 90 min at 32°C) in the presence of Polybrene (5 μg/ml) on the plates coated with Retronectin (50 μg/ml) (Takara/Clontech) and anti-CD3 (1-2 μg/ml). Human Jurkat T cells were directly transduced with lentiviral particles by spinfection. Five days after transduction, cells were sorted on the basis of expression of GFP. GFP expressing human Jurkat T cells were further purified by FACS at least three times before using for experiments.

### Proximity ligation assay (PLA)

PLA was performed with Duolink In Situ Detection Reagents Orange (Sigma) according to the manufacturer’s recommendation with minor modifications. Tregs were cultured for four days and harvested, and cells were fixed with 2% paraformaldehyde for 10 min at RT. Fixed cells were incubated in Foxp3 Fix/Perm buffer set for 30 min at 4°C, followed by staining with mouse anti-β-catenin and rabbit anti-Foxo1 for 1 h at RT in Foxp3 staining buffer. Cells were washed and stained in Foxp3 staining buffer with the secondary mouse PLUS and rabbit MINUS antibodies for 30 min at RT. Cells were washed in TBS (0.01 M Tris, 0.15 M NaCl) with 0.5% BSA and the ligation reaction was performed at 37°C for 30 min, followed by the amplification reaction at 37°C for 100 min. Cells were washed in TBS (0.2 M Tris, 0.1 M NaCl) with 0.5% BSA and stained with anti-Foxp3 antibody for 30 min at 4°C. Cells were analyzed with a 60x or 100x objective on a Leica DM6000 CS confocal microscope.

### Statistical analysis

All statistical analyses were performed using GraphPad Prism 6 (GraphPad Software). Detailed information about statistical analysis, including tests and values used, is provided in the figure legends. Values of *P*<0.05 or less were considered significant.

## Data availability statement

The data that support the findings of this study are available from the corresponding authors upon request.

## Acknowledgements

We thank L. Devine, C. Wang, and H. Tomita for cell sorting, and Ee-chun Cheng and M. Zhang for preparation of the RNA-seq library, K. Tanaka for microscopic imaging, and M. Shimizu, H. Taniwaki, and N. Yamanaka for technical support. FIC mice were a generous gift from S. Sakaguchi (Osaka University). This work was supported by grants from the Uehara Memorial Foundation Research Fellowship, MSD Life Science Foundation Research Fellowship, LEGEND Study Abroad Grant from BioLegend and Tomy Digital Biology (to T.S.), the Ministry of Education, Culture, Sports, Science and Technology (MEXT); JSPS KAKENHI (Grant Number 21229010), the Core Research for Evolutional Science and Technology (CREST) program from the Japan Science and Technology Agency (to I.K.), National Institutes of Health Grants (P01 AI045757, U19 AI046130, U19 AI070352, and P01 AI039671), and the Nancy Taylor Foundation for Chronic Diseases (to D.A.H.). RNA sequencing service was conducted at Yale Stem Cell Center Genomics Core facility that was supported by the Connecticut Regenerative Medicine Research Fund and the Li Ka Shing Foundation.

## Contributions

T.S., D.M.R., and C.M.U. performed *in vitro* experiments with the help of M.R.L. and M.D.V.; T.S. and A.T.N. performed *in vivo* experiments with the help of H.A. and T.N.; T.S. and M.R.L. analyzed the RNA-seq data under the supervision of D.A.H.; T.S. performed data analysis and wrote the manuscript under the supervision of A.T.N., I.K., M.R.L., M.D.V., and D.A.H.; A.T.N, I.K., M.D.V. and D.A.H. supervised the overall study.

## Competing interests

The authors declare no competing financial interests.

## Corresponding author

Correspondence to Tomokazu Sumida.

## Supplementary Figure legends

**Supplementary Figure 1.**
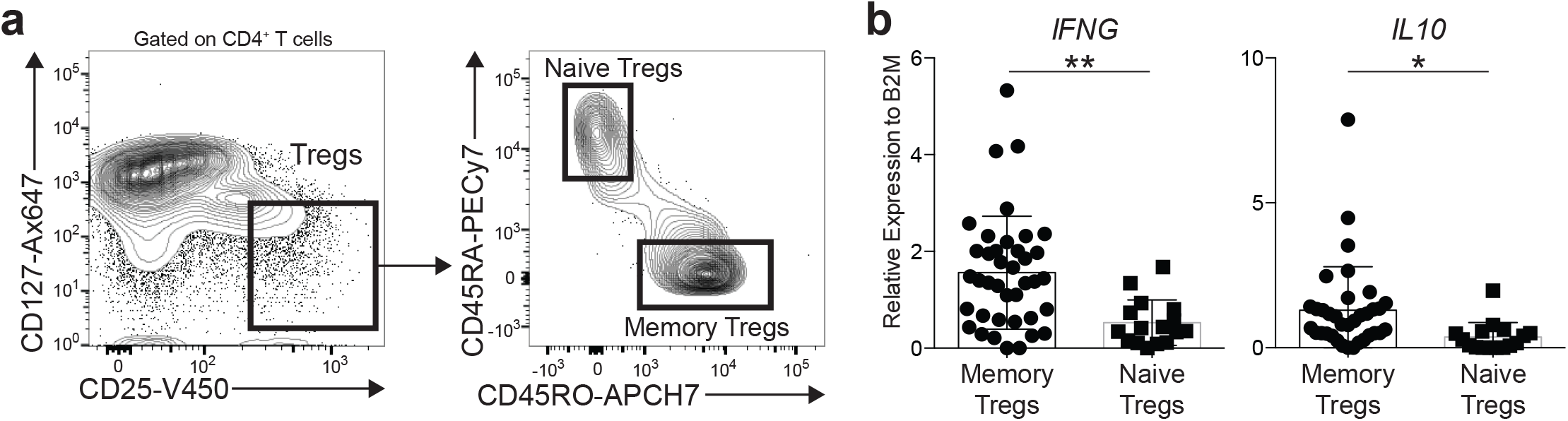
Memory Tregs are the main source of effector cytokines IFNγ and IL-10. **(a)** Sorting strategy for memory and naive Tregs from circulating human CD4^+^ T cells. **(b)** mRNA expression of *IFNG* and *IL10* gene on memory and naive Tregs (memory Tregs; n=35 subjects, naïve Tregs; n=16 subjects). \**P*<0.05, \*\**P*<0.01 (two-tailed unpaired Student’s *t*-test).

**Supplementary Figure 2.**
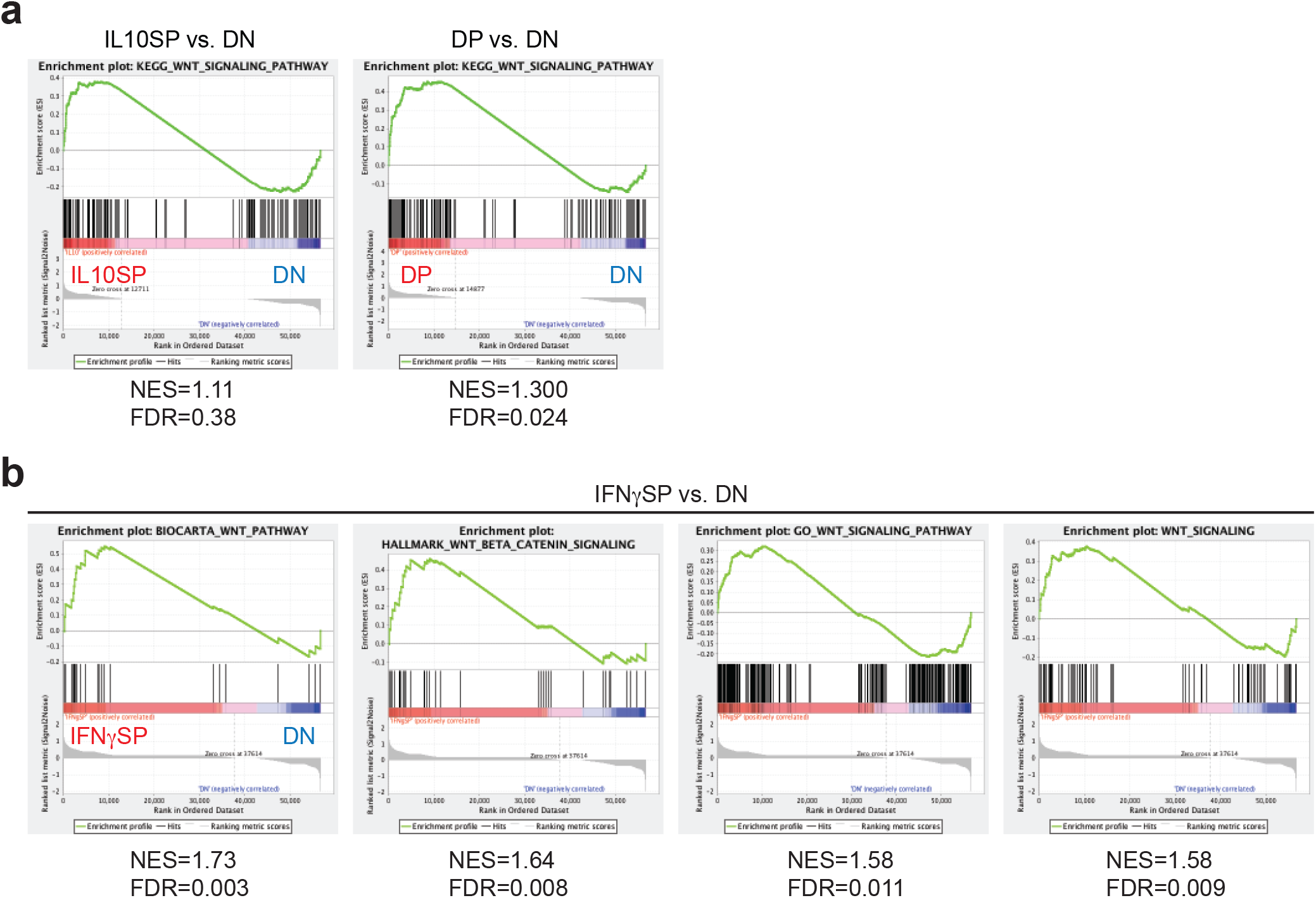
GSEA of Wnt signaling pathway on Treg subsets. **(a)** GSEA enrichment plots of KEGG Wnt signaling pathway between IL10SP vs. DN and DP vs. DN. **(b)** GSEA enrichment plots of four different Wnt signaling pathway gene sets between IFNγSP vs. DN.

**Supplementary Figure 3.**
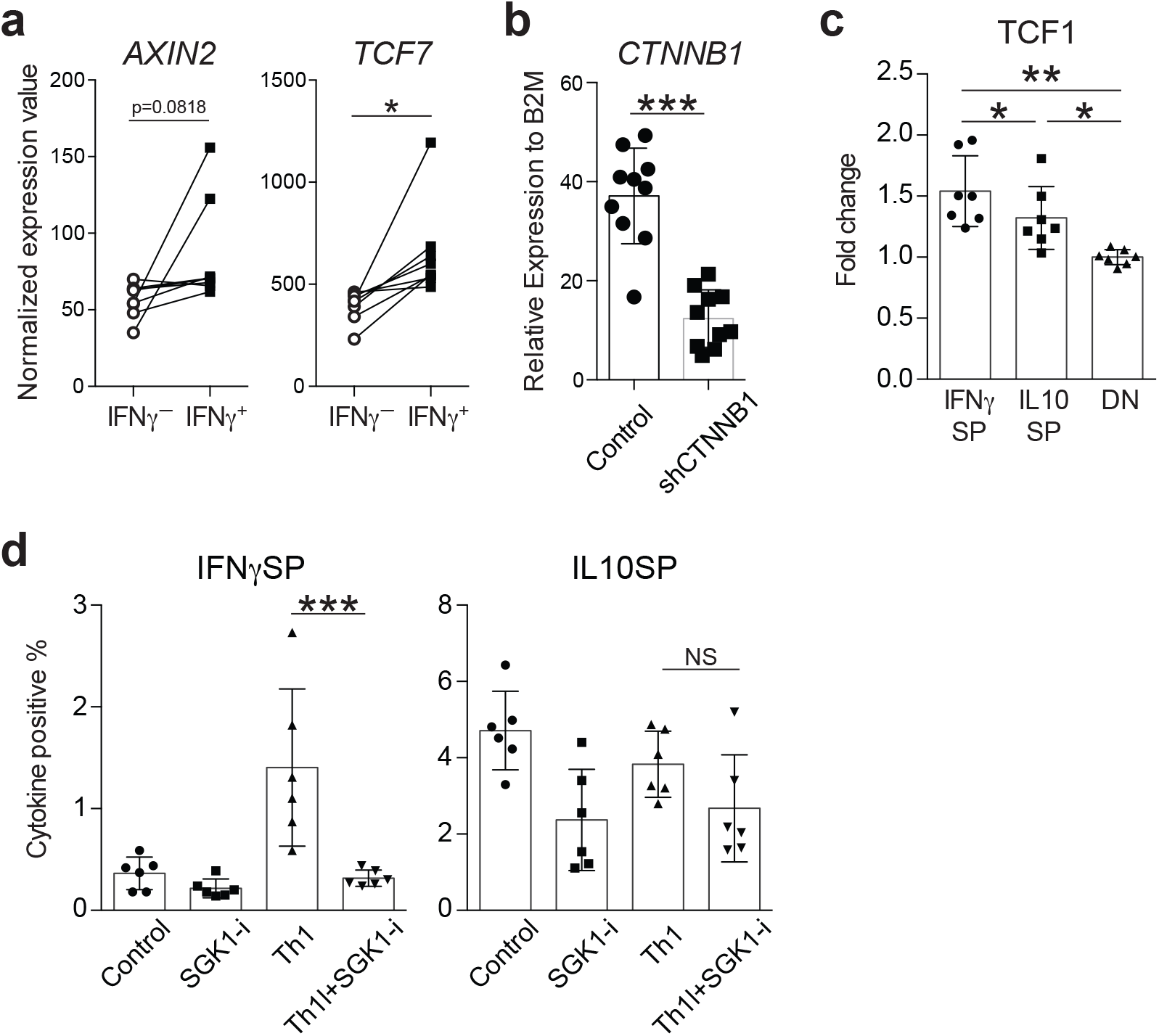
β-catenin signaling is activated in IFNγ producing human Treg subset. **(a)** *AXIN2* and *TCF7* mRNA expression in IFNγ^+^ and IFNγ^−^ human Treg populations assessed by DNA microarray (n=8 subjects)^32^. \**P*<0.05 (two-tailed paired Student’s *t*-test). **(b)** *CTNNB1* gene expression on Tregs transduced with a non-targeted shRNA or a *CTNNB1* shRNA and cultured for 5 days (n=10 subjects). \*\*\**P*<0.001 (two-tailed unpaired Student’s *t*-test). **(c)** Flow cytometric analysis of TCF1 expression on *ex vivo* Treg subpopulations relative to DN (n=8 subjects). \**P*<0.05, \*\**P*<0.01 (one-way ANOVA with Tukey’s multiple comparisons test). **(d)** Frequency of IFNγ and IL-10 positive cell number. Tregs were stimulated with anti-CD3 and anti-CD28 in the presence of SGK1 inhibitor GSK650394 (SGK1-i), IL-12 (Th1), or IL-12 and GSK650394 (Th1^+^ SGK1-i) (n=6 subjects) \*\*\**P*<0.001 (one-way ANOVA with Tukey’s multiple comparisons test).

**Supplementary Figure 4.**
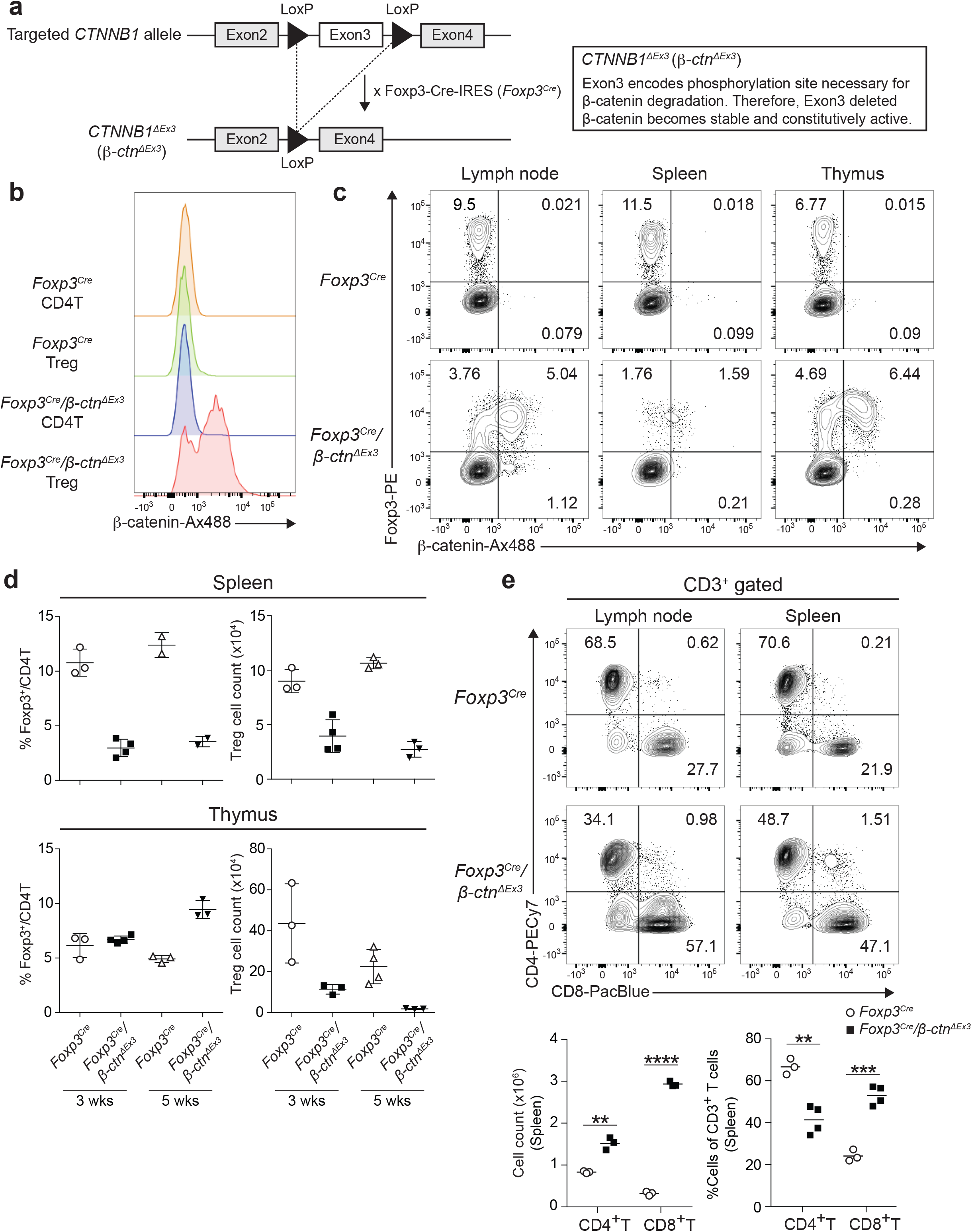
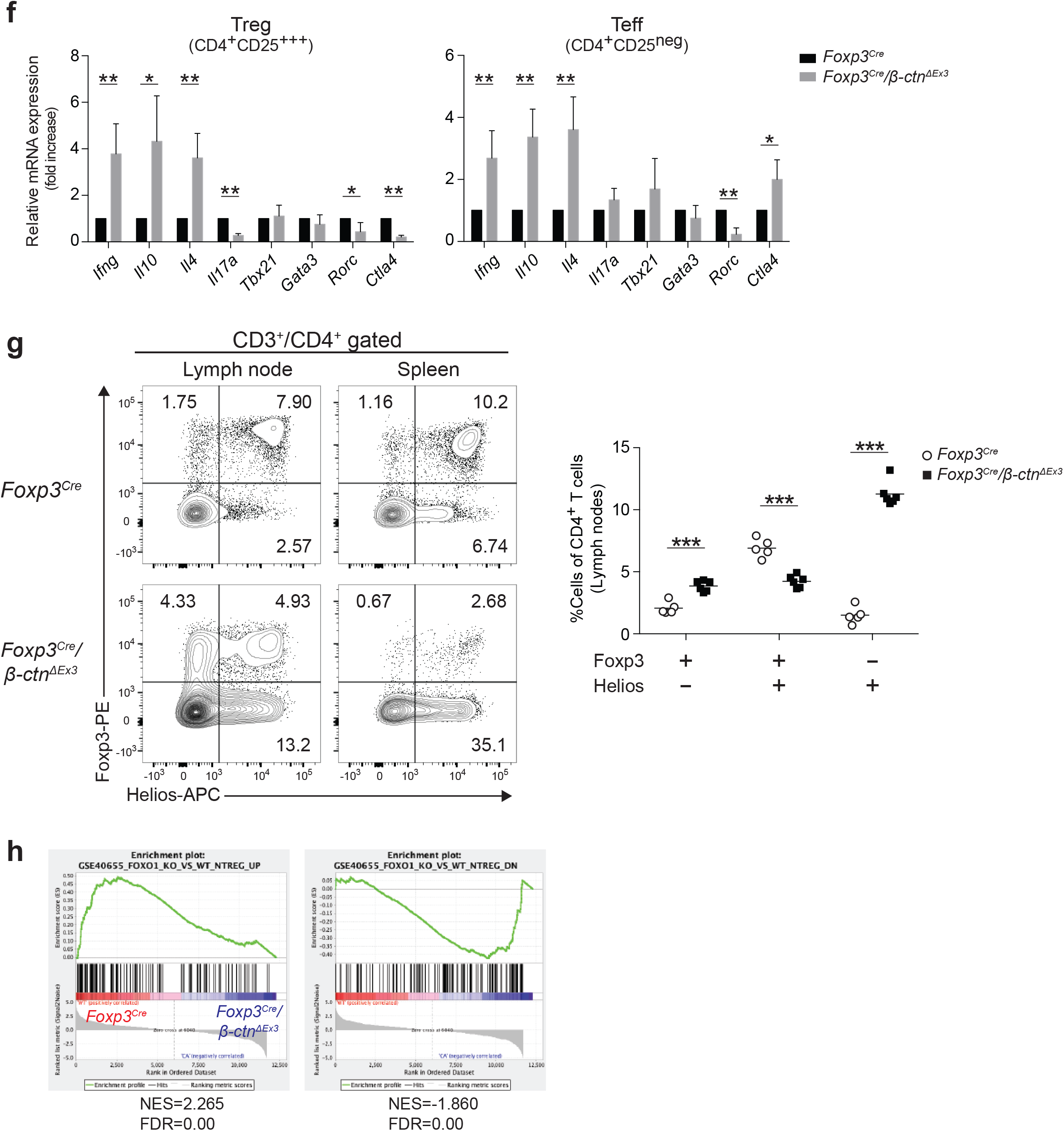
Treg specific activation of β-catenin induces IFNγ secreting dysfunctional phenotype with *Scurfy*-like autoimmunity. **(a)** Schematic of the wild-type and targeted *CTNNB1* allele. Exon3 of *CTNNB1* gene encodes a phosphorylation site necessary for β-catenin degradation. Therefore, Exon3 deleted β-catenin can escaped degradation and works as a constitutively active form. **(b)** Flow cytometric analysis of β-catenin on peripheral lymph node Foxp3^+^ Tregs (Treg) and Foxp3^−^ CD4^+^ T cells (CD4T) from *Foxp3^Cre^* and *Foxp3^Cre^/β-ctn^ΔEx3^* mice. **(c)** Flow cytometric analysis of β-catenin and Foxp3 in peripheral lymph nodes, spleen, and thymus CD4^+^ T cells from *Foxp3^Cre^* and *Foxp3^Cre^/β-ctn^ΔEx3^* mice. **(d)** The percentage of Foxp3^+^ Tregs within CD4^+^ T cells and the cell numbers of Foxp3^+^ Tregs in spleen (top) and thymus (bottom) from *Foxp3^Cre^* and *Foxp3^Cre^/β-ctn^ΔEx3^* mice at 3 weeks (3wks) and 5 week old (5wks) of age. **(e)** Flow cytometric analysis of CD4^+^ and CD8^+^ T cells in peripheral lymph nodes and spleen from *Foxp3^Cre^* and *Foxp3^Cre^/β-ctn^ΔEx3^* mice at the age of 3 weeks. Cell count and percentages of CD4^+^ and CD8^+^ T cells among CD3^+^ T cells from the spleen were shown at the bottom. \*\**P*<0.01 \*\*\**P*<0.001, \*\*\*\**P*<0.0001 (two-way ANOVA with Sidak’s multiple comparisons test). **(f)** Expression for classical helper cytokines and transcription factors in both Tregs and T effector cells (CD4^+^ CD25^neg^) assessed by qPCR (n=4 mice). \**P*<0.05, \*\**P*<0.01 (two-tailed Student’s *t*-test). **(g)** Flow cytometric analysis of Foxp3 and Helios expression on CD4^+^ T cells in peripheral lymph nodes and spleen from *Foxp3^Cre^* and *Foxp3^Cre^/β-ctn^ΔEx3^* mice at the age of 3 weeks. Percentages of Foxp3^+^ and/or Helios+ CD4^+^ T cells isolated from lymph nodes are shown at the bottom. (n=5-6 mice) \*\*\**P*<0.001 (two-way ANOVA with Sidak’s multiple comparisons test). **(h)** GSEA enrichment plot between *Foxp3^Cre^* and *Foxp3^Cre^/β-ctn^ΔEx3^* Tregs using the gene set that is positively regulated by Foxo1 (left) and negatively regulated by Foxo1 (right) identified from the comparison between Wild type vs. Foxo1 KO Tregs (GSE40655). Normalized enrichment score (NES) and false discovery rate (FDR) are indicated.

**Supplementary Figure 5.**
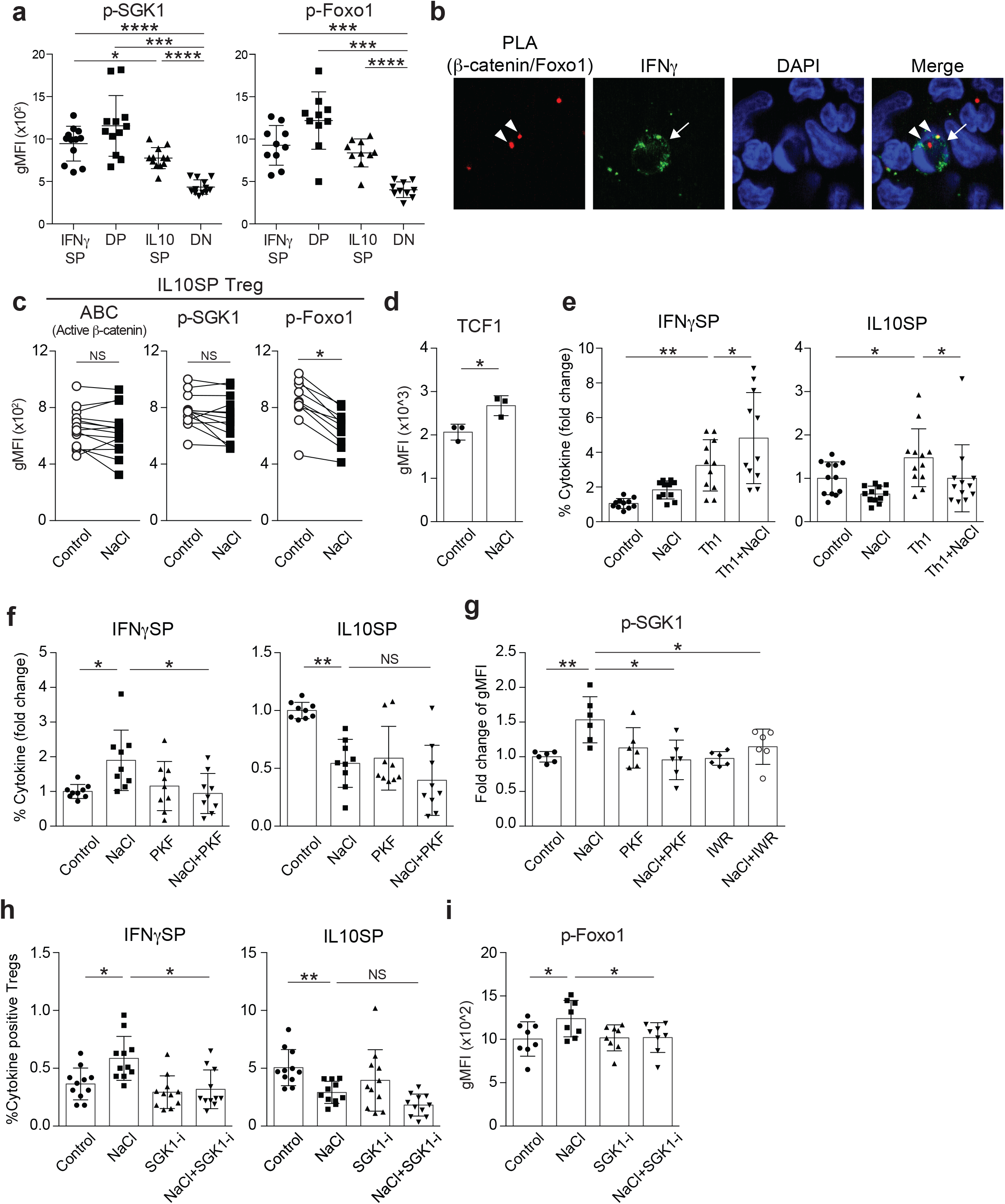
High salt activates the β-catenin/SGK1/Foxo axis in IFNγ-producing human Tregs. **(a)** Flow cytometric analysis of phospho-SGK1 (left) and phospho-Foxo1 (right) level in human Treg subsets. Tregs were stimulated with anti-CD3 and anti-CD28 for 96 h followed by 4 h PMA/iomomycin stimulation, and the expression of p-SGK1 and p-Foxo1 were determined by intracellular staining in each subset. (n=12 subjects; pSGK1, n=10 subjects; p-Foxo1) **(b)** Representative immunofluorescence images of human Tregs with PLA signal for β-catenin-Foxo1 interaction (red) and IFNγ staining (green). Nuclei were stained with DAPI (blue). PLA signal (arrowheads) was observed in an IFNγ positive cell (arrow). **(c)** Flow cytometric analysis of ABC, phospho-Foxo1 (Ser256), and phospho-SGK1 (Thr256) expression in human IL-10 producing Tregs. Tregs were stimulated with anti-CD3 and anti-CD28 in the presence (NaCl) or absence (Control) of additional 40 mM NaCl for 96 h followed by PMA/iomomycin stimulation for 4 h (n=10-15 subjects). \**P*<0.05 (two-tailed unpaired Student’s *t*-test). **(d)** Flow cytometric analysis of TCF1 expression in human IFNγ producing Tregs. Tregs were cultured as in **(c)** (n=3 subjects). \**P*<0.05 (two-tailed unpaired Student’s *t*-test). **(e)** Relative frequency of IFNγ and IL-10 positive cell number (fold of control condition) in human Tregs cultured as in Fig. 5c (n=11 subjects). \**P*<0.05, \*\**P*<0.01 (one-way ANOVA with Tukey’s multiple comparisons test). **(f)** Relative frequency of IFNγ and IL-10 positive cell number (fold of control condition) in human Tregs cultured as in Fig. 5d (n=9 subjects). \**P*<0.05, \*\**P*<0.01 (one-way ANOVA with Tukey’s multiple comparisons test). **(g)** Flow cytometric analysis of p-SGK1 expression in human IFNγ producing Tregs. Tregs were cultured as in Fig. 5d (n=6 subjects). \**P*<0.05, \*\**P*<0.01 (one-way ANOVA with Tukey’s multiple comparisons test). **(h)** Frequency of IFNγ and IL-10 positive cell number in human Tregs stimulated in the presence (NaCl) or absence (Control) of additional 40 mM NaCl with and without Wnt/inhibitor GSK650394 (SGK1-i) for 96 h (n=11 subjects). \**P*<0.05, \*\**P*<0.01 (one-way ANOVA with Tukey’s multiple comparisons test). **(i)** Flow cytometric analysis of p-Foxo1 expression in human IFNγ producing Tregs. Tregs were cultured as in **(h)** (n=8 subjects). \**P*<0.05 (one-way ANOVA with Tukey’s multiple comparisons test).

**Supplementary Figure 6.**
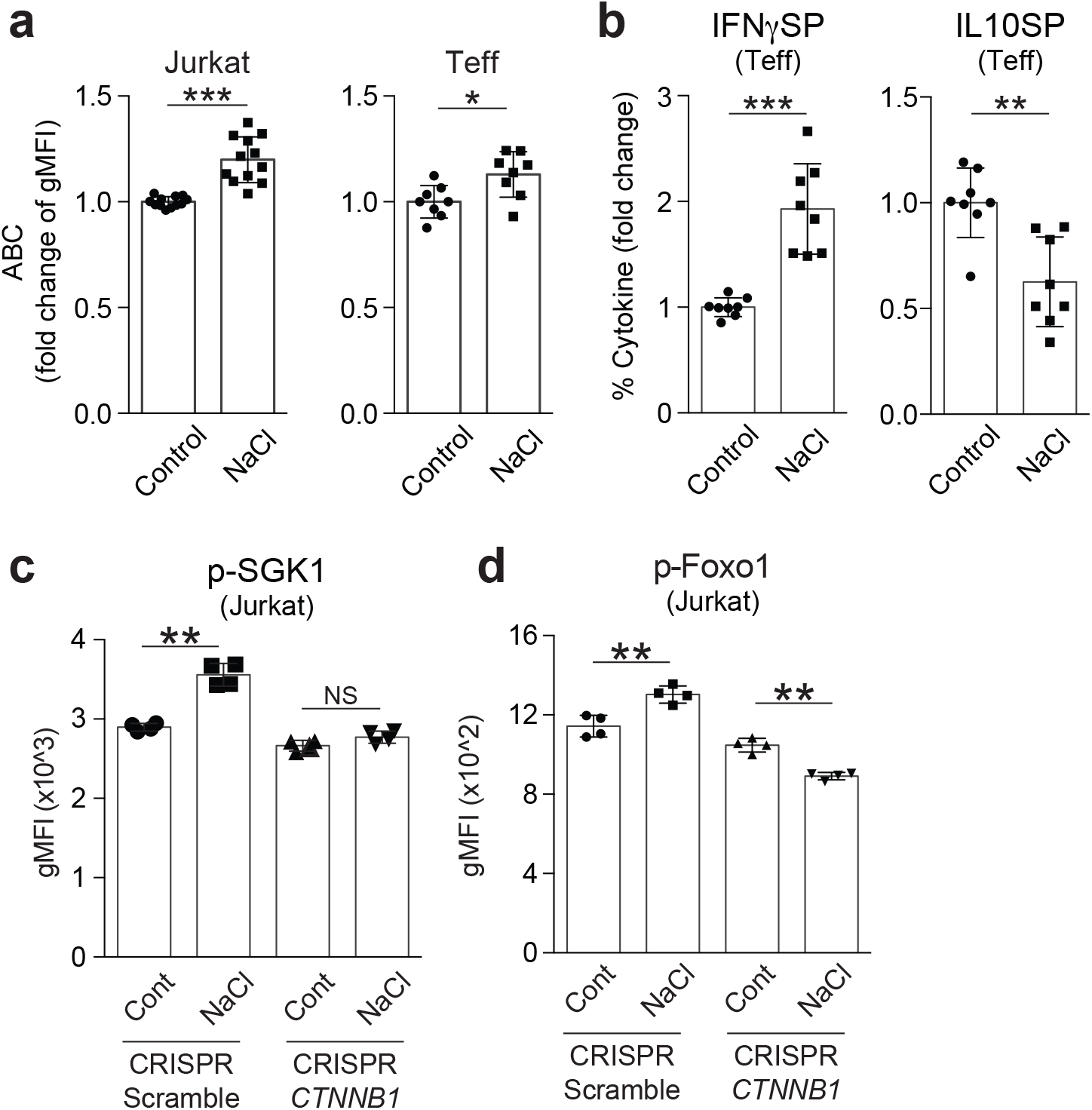
The β-catenin/SGK1/Foxo axis is also activated in Teff and human Jurkat T cells under high salt conditions. **(a)** Flow cytometric analysis of ABC expression in human T effector cells (Teffs) and human Jurkat T cells. Human Teff were stimulated as well as Tregs for 96 h (n=8 subjects). Human Jurkat T cells were cultured without TCR stimulation for 120 h (n=12). Both were cultured in the presence (NaCl) or absence (Control) of additional 40 mM NaCl. \**P*<0.05, \*\*\**P*<0.001 (two-tailed unpaired Student’s *t*-test). **(b)** Relative frequency of IFNγ and IL-10 positive cell number (fold of control condition) in human Teffs cultured as in **(a)** (n=8 subjects). \*\**P*<0.01, \*\*\**P*<0.001 (two-tailed unpaired Student’s *t*-test). **(c, d)** Flow cytometric analysis of SGK1 phosphorylation at Thr256 **(c)** and Foxo1 phosphorylation at Ser256 **(d)** in human Jurkat T cells. Human Jurkat T cells were transduced with scramble gRNA (CRISPR/Scramble) or *CTNNB1* targeted gRNA (CRISPR/CTNNB1) with Cas9. Both cell lines were cultured in the presence (NaCl) or absence (Control) of additional 40 mM NaCl without TCR stimulation for 120 h (n=4). \*\**P*<0.01 (one-way ANOVA with Tukey’s multiple comparisons test).

**Supplementary Figure 7.**
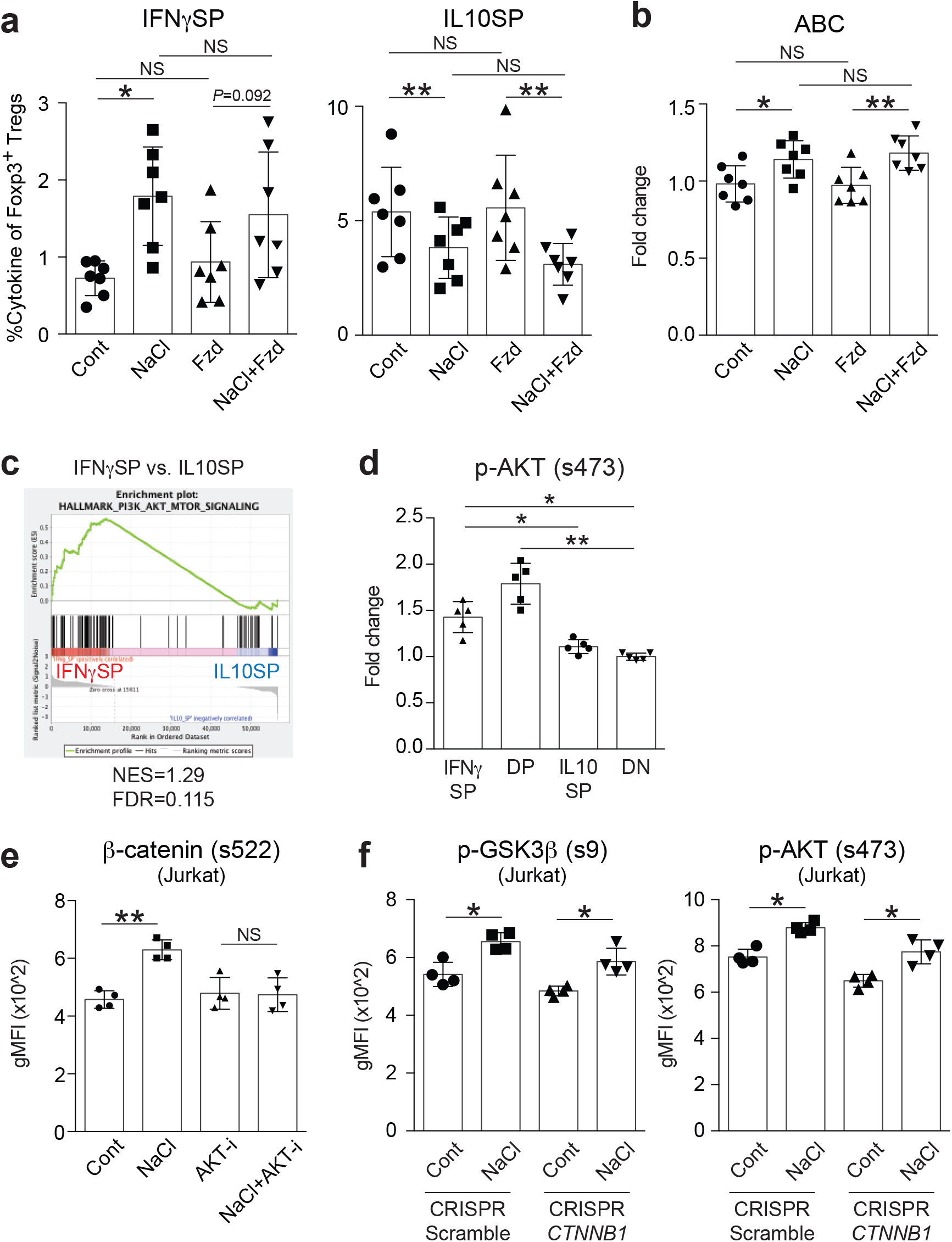
High salt induced β-catenin activation via AKT is independent of Wnt ligands. **(a)** Relative frequency of IFNγ and IL-10 positive cell number (fold of control condition) in human Tregs. Tregs were stimulated with anti-CD3 and anti-CD28 in the presence of Fzd8-FC (Fzd), additional 40 mM NaCl (NaCl), or Fzd8-FC and NaCl (NaCl + Fzd8) (n=7 subjects). \**P*<0.05, \*\**P*<0.01 (one-way ANOVA with Tukey’s multiple comparisons test). **(b)** Relative expression level of ABC in human Tregs cultured as in **(a)**. (n=7 subjects) \**P*<0.05, \*\**P*<0.01 (one-way ANOVA with Tukey’s multiple comparisons test). **(c)** GSEA enrichment plots of PI3K/AKT pathway gene sets between IFNγSP vs. IL10SP. Normalized enrichment score (NES) and false discovery rate (FDR) are indicated at the bottom of each plot. **(d)** Relative expression level of p-AKT on Treg subsets. Tregs were stimulated with anti-CD3 and anti-CD28 for 4 days, followed by 4 h PMA/iomomycin stimulation and intracellular cytokine staining for IFNγ and IL-10 (n=5 subjects). \**P*<0.05, \*\**P*<0.01 (one-way ANOVA with Tukey’s multiple comparisons test). **(e)** Flow cytometric analysis of β-catenin phosphorylation at s522 in human Jurkat T cells. Human Jurkat T cells were stimulated in the presence of AKT inhibitor MK2206 (AKT-i), additional 40 mM NaCl (NaCl), or MK2206 and NaCl (NaCl + AKT-i) (n=4). **(f)** Flow cytometric analysis of GSK3β phosphorylation at s9 and AKT phosphorylation at s473 in human Jurkat T cells. Human Jurkat T cells were prepared as in Supplementary Fig. 6c. (n=4). \**P*<0.05, \*\**P*<0.01 (one-way ANOVA with Tukey’s multiple comparisons test).

**Supplementary Figure 8.**
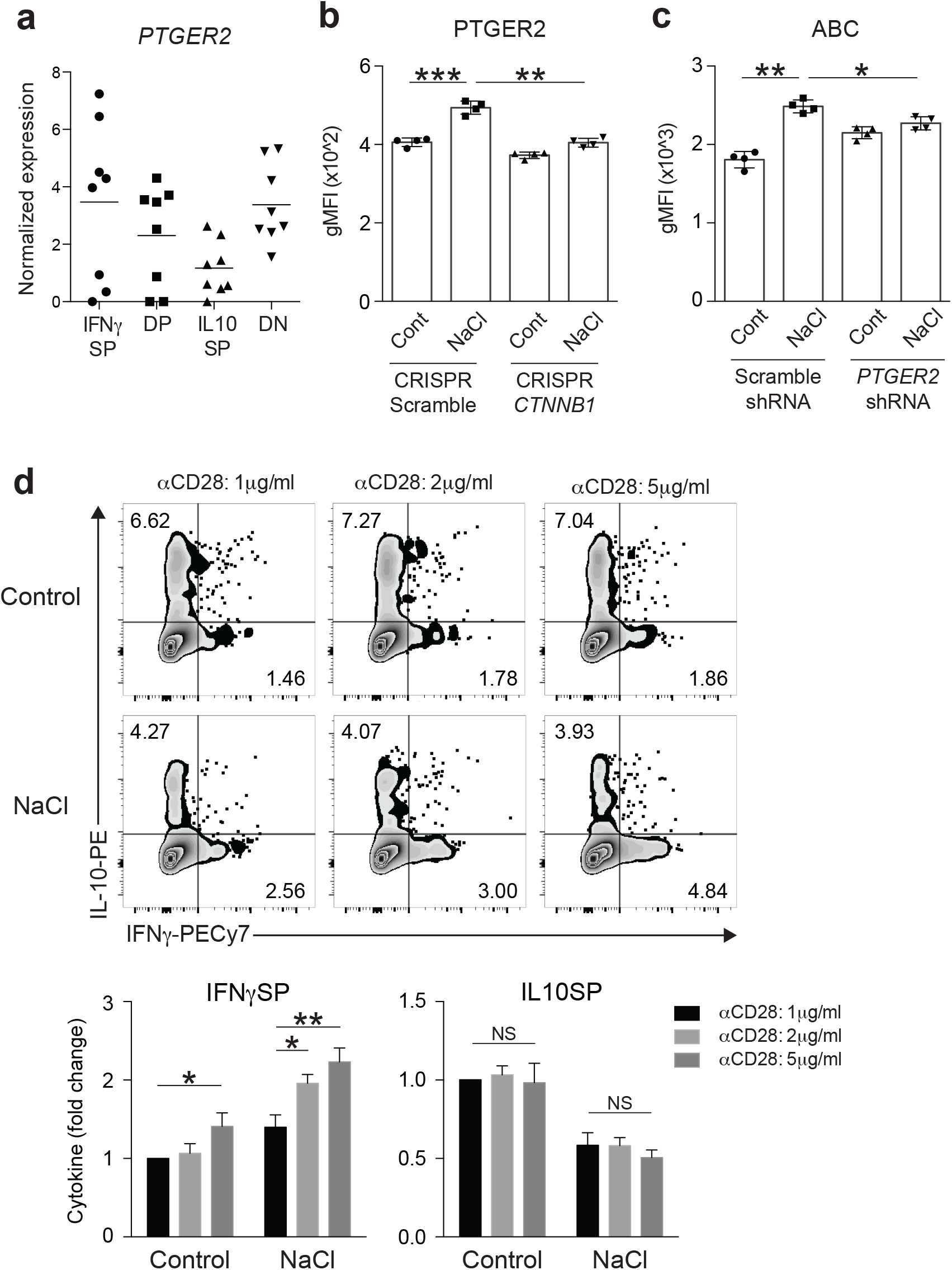
PTGER2-β-catenin loop is activated by high salt stimulation. **(a)** *PTGER2* expression assessed by RNA-seq on *ex vivo* Treg subpopulations (n=8 subjects). **(b)** Flow cytometric analysis of PTGER2 in human Jurkat T cells. Human Jurkat T cells were prepared as in Supplementary Fig. 6c. (n=4). \*\**P*<0.01, \*\*\**P*<0.001 (one-way ANOVA with Tukey’s multiple comparisons test). **(c)** Flow cytometric analysis of ABC in human Jurkat T cells. Human Jurkat T cells were transduced with a scramble shRNA or a *PTGER2* shRNA and cultured in normal media (Control) or media supplemented with additional 40 mM NaCl (NaCl) for 120 h. (n=4) \**P*<0.05, \*\**P*<0.01 (one-way ANOVA with Tukey’s multiple comparisons test). **(d)** Representative flow cytometric analysis of IFNγ and IL-10 production in human Tregs cultured in the normal media (Control) or media supplemented with additional 40 mM NaCl (NaCl) with anti CD3 (2μg/ml) and different concentration of anti CD28 (1, 2, 5 μg/ml) for 96 h. Relative frequency of IFNγ and IL-10 producing Tregs are shown at the bottom (n=4 subjects). \**P*<0.05, \*\**P*<0.01 (two-way ANOVA with Sidak’s multiple comparisons test).

**Supplementary Figure 9.**
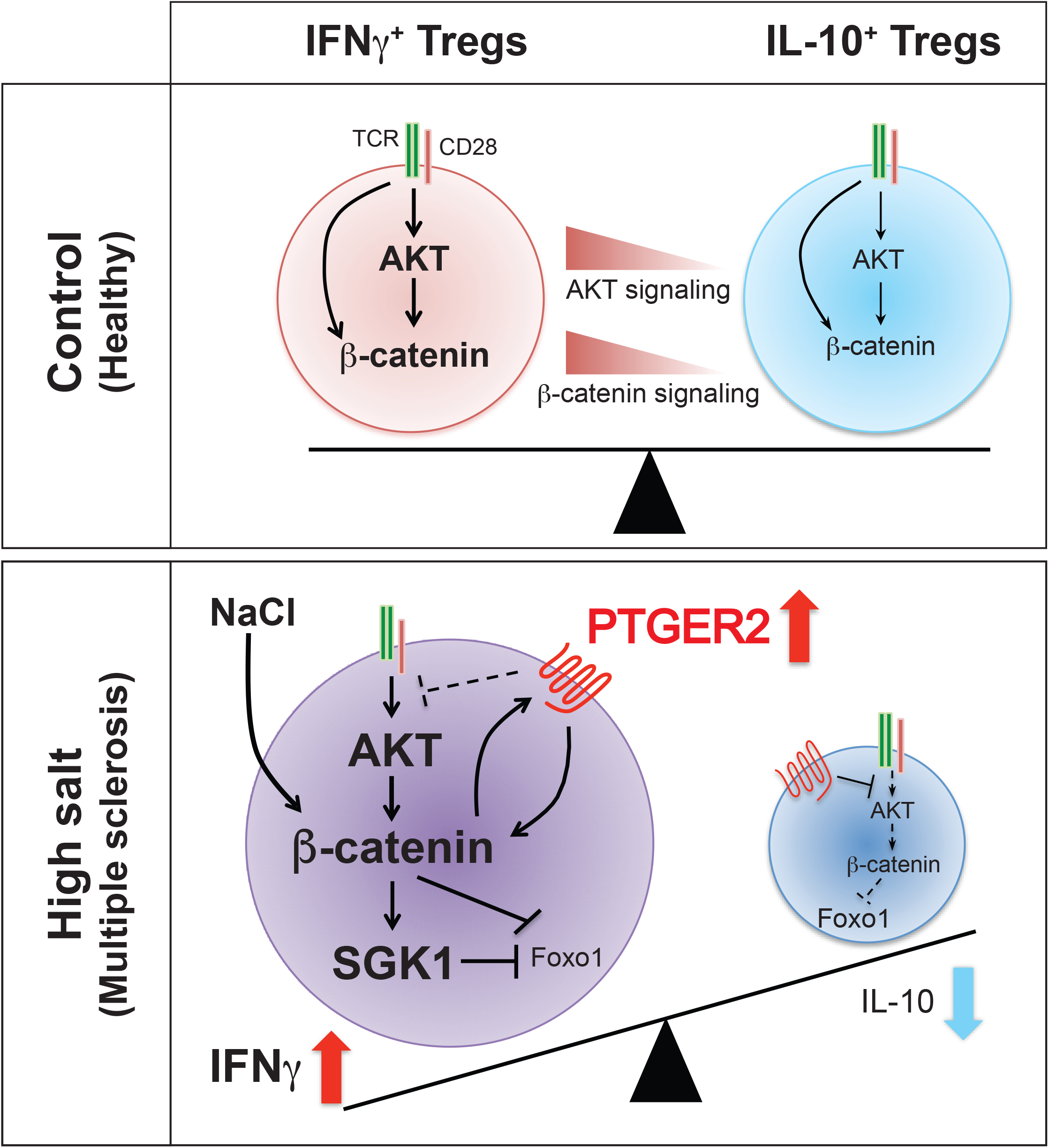
Schematic model of the role of PTGER2 and the AKT/β-catenin/SGK1/Foxo axis for the production of IFNγ and IL-10 in Tregs. AKT/β-catenin signaling balances IFNγ/IL-10 production in Tregs. Under high salt conditions, PTGER2 was increased and established the positive feed forward loop with β-catenin, resulted in amplified activation of the β-catenin/SGK1/Foxo axis in IFNγ-producing Tregs.

Supplementary Table 1

Supplementary Table 1
| Patients | Age (at time of collection) | Sex | Ethnicity | Duration of Disease | Treatment | EDSS | IFNG mRNA expression | IL10 mRNA expression | Ratio of IFNG/IL10 mRNA expression |
| --- | --- | --- | --- | --- | --- | --- | --- | --- | --- |
| #1 | 57 | M | African american | 16 yrs | untreated | 4.5 | 0.34 | 1.98 | 0.172 |
| #2 | 46 | F | Caucasian/Non-Hispanic | 7 yrs | untreated | 2.5 | 1.54 | 4.83 | 0.319 |
| #3 | 36 | F | Caucasian/Hispanic | 1 mo | untreated | 3.5 | 0.55 | 21.44 | 0.026 |
| #4 | 29 | F | Caucasian/Non-Hispanic | 3 yrs | untreated | 2 | 1.31 | 6.29 | 0.208 |
| #5 | 48 | F | Caucasian/Non-Hispanic | 18 yrs | untreated | 6.5 | 0.72 | 3.07 | 0.235 |
| #6 | 41 | M | Caucasian/Non-Hispanic | 10 yrs | untreated | 1.5 | 2.32 | 9.37 | 0.248 |
| #7 | 31 | M | Caucasian/Non-Hispanic | 2 mo | untreated | 6 | 0.37 | 3.78 | 0.098 |
| #8 | 41 | F | Caucasian/Non-Hispanic | 3 mo | untreated | 3 | 0.09 | 24.75 | 0.004 |
| #9 | 42 | M | Caucasian/Non-Hispanic | <1 mo | untreated | 1.5 | 0.4 | 11.63 | 0.034 |
| #10 | 34 | F | Caucasian/Non-Hispanic | 4 mo | untreated | 4 | 0.57 | 2.97 | 0.192 |
| #11 | 38 | M | Caucasian/Non-Hispanic | <1 mo | untreated | 0 | 0.35 | 4.23 | 0.083 |
| #12 | 43 | F | African american | 1 yr | untreated | 1 | 0.24 | 3 | 0.080 |
| #13 | 53 | F | Caucasian/Non-Hispanic | 28 yrs | untreated | 4.5 | 1.27 | 1.34 | 0.948 |
| #14 | 32 | F | Caucasian/Non-Hispanic | 1 mo | untreated | 1.5 | 0.99 | 18.9 | 0.052 |
| #15 | 32 | F | Caucasian/Non-Hispanic | 3 yrs | untreated | not assessed | 0.34 | 3.15 | 0.108 |
| #16 | 28 | F | Caucasian/Non-Hispanic | 2 yrs | untreated | 1 | 0.43 | 7.69 | 0.056 |
| #17 | 50 | F | Caucasian/Non-Hispanic | <1 mo | untreated | 3.5 | 1.81 | 12.58 | 0.144 |
| #18 | 43 | F | Caucasian/Non-Hispanic | 5 mo | untreated | 2.5 | 2.1 | 3.64 | 0.577 |
| #19 | 23 | F | Caucasian/Non-Hispanic | <1 mo | untreated | 1 | 1.08 | 4.3 | 0.251 |
| #20 | 58 | F | Caucasian/Non-Hispanic | 13 yrs | untreated | 6 | 2.58 | 2.23 | 1.157 |
| #21 | 34 | F | Caucasian/Non-Hispanic | 1 mo | untreated | 1.5 | 2.1 | 9.81 | 0.214 |
| #22 | 30 | F | Caucasian/Non-Hispanic | 12 yrs | untreated | 2 | 2.47 | 30.16 | 0.082 |
| #23 | 27 | F | Caucasian/Non-Hispanic | <1 mo | untreated | 0 | 1.78 | 13.38 | 0.133 |
| #24 | 61 | F | Caucasian/Non-Hispanic | 20 yrs | untreated | 4 | 1.05 | 12.99 | 0.081 |
| #25 | 35 | F | Caucasian/Hispanic | 1 yr | untreated | 2 | 0.89 | 21.71 | 0.041 |
| #26 | 33 | F | Caucasian/Non-Hispanic | 1 mo | untreated | 1.5 | 0.5 | 18.97 | 0.026 |
| #27 | 44 | F | Caucasian/Non-Hispanic | 8 yrs | untreated | 2.5 | 0.46 | 14.56 | 0.032 |

Supplementary Table 2

Supplementary Table 2
| *Antibodies/Kits* | *Source* | *Experiments for use* |
| --- | --- | --- |
| **Human** |  |  |
| anti-CD4 (RPA-T4) | BD Bioscience | Flow cytometry |
| anti-CD25 (MA251) | BD Bioscience | Flow cytometry |
| anti-CD127 (HIL-7R-M21) | BD Bioscience | Flow cytometry |
| anti-CD45RO (UCHL1) | BD Bioscience | Flow cytometry |
| anti-IL-10 (JES3-9D7) | BioLegend | Flow cytometry |
| anti-IFNγ (B27) | BD Bioscience | Flow cytometry |
| anti-IFNγ (4S.B3) | eBioscience | Flow cytometry |
| anti-β-catenin (14/Beta-Catenin) | BD Bioscience | Flow cytometry, PLA |
| anti-active β-catenin (8E7) | Millipore | Flow cytometry |
| anti-phospho β-catenin (Ser522) (D8E11) | CST | Flow cytometry |
| anti-phospho AKT (Ser473) (D9E) | CST | Flow cytometry |
| anti-phospho GSK3β (Ser9) (D85E12) | CST | Flow cytometry |
| anti-Foxp3 (PCH101) | eBioscience | Flow cytometry |
| anti-phospho Foxo1 (S256) polyclonal | Bioss | Flow cytometry |
| anti-phospho SGK1 (T256) polyclonal | Bioss | Flow cytometry |
| IFNγ secretion assay (APC) | Miltenyi | Flow cytometry |
| IL-10 secretion assay (PE) | Miltenyi | Flow cytometry |
| anti-Foxo1 (C29H4) | CST | PLA |
| anti-TCF1 (S33-966) | BD Bioscience | Flow cytometry |
| anti-Tbet (4B10) | BioLegend | Flow cytometry |
| anti-PTGER2 (EPR8030(B)) | Abcam | Flow cytometry |
| anti-CD3 (UCHT1) | BD Bioscience | Cell culture |
| anti-CD28 (28.2) | BD Bioscience | Cell culture |
| **Mouse** |  |  |
| anti-Foxp3 (FJK-16s) | eBioscience | Flow cytometry |
| anti-CD3 (145-2C11) | eBioscience | Flow cytometry |
| anti-CD4 (RM4-5) | eBioscience | Flow cytometry |
| anti-CD8 (53-6.7) | eBioscience | Flow cytometry |
| anti-Helios (22F6) | BioLegend | Flow cytometry |
| anti-phospho Foxo1 (S256) (E1F7T) | CST | Flow cytometry |
| anti-phospho Foxo3a (Ser253) (D18H8) | CST | Flow cytometry |
| anti-phospho Foxo1(T24)/3a(T32)/4(T28) (4G6) | CST | Flow cytometry |
| anti-SGK1 (Y238) | Abcam | Flow cytometry |
| anti-GATA3 (TWAJ) | eBioscience | Flow cytometry |
| anti-RORγt (B2D) | eBioscience | Flow cytometry |
| Dynabeads Mouse T-Activator CD3/CD28 | Invitrogen | Cell culture |
| **Human/Mouse** |  |  |
| PE-Cy™7 Streptavidin | BD Bioscience | Flow cytometry |
| Zombie Aqua™ Fixable Viability dye | BioLegend | Flow cytometry |

## References

1. Sakaguchi S, Ono M, Setoguchi R, Yagi H, Hori S, Fehervari Z, et al. Foxp3+ CD25+ CD4+ natural regulatory T cells in dominant self-tolerance and autoimmune disease. Immunol Rev 2006, 212: 8–27.

2. Bennett CL, Christie J, Ramsdell F, Brunkow ME, Ferguson PJ, Whitesell L, et al. The immune dysregulation, polyendocrinopathy, enteropathy, X-linked syndrome (IPEX) is caused by mutations of FOXP3. Nat Genet 2001, 27(1): 20–21.

3. Lahl K, Loddenkemper C, Drouin C, Freyer J, Arnason J, Eberl G, et al. Selective depletion of Foxp3+ regulatory T cells induces a scurfy-like disease. J Exp Med 2007, 204(1): 57–63.

4. Viglietta V, Baecher-Allan C, Weiner HL, Hafler DA. Loss of functional suppression by CD4+CD25+ regulatory T cells in patients with multiple sclerosis. J Exp Med 2004, 199(7): 971–979.

5. Buckner JH. Mechanisms of impaired regulation by CD4(+)CD25(+)FOXP3(+) regulatory T cells in human autoimmune diseases. Nat Rev Immunol 2010, 10(12): 849–859.

6. Vignali DA, Collison LW, Workman CJ. How regulatory T cells work. Nat Rev Immunol 2008, 8(7): 523–532.

7. Miyara M, Yoshioka Y, Kitoh A, Shima T, Wing K, Niwa A, et al. Functional delineation and differentiation dynamics of human CD4+ T cells expressing the FoxP3 transcription factor. Immunity 2009, 30(6): 899–911.

8. Dominguez-Villar M, Baecher-Allan CM, Hafler DA. Identification of T helper type 1-like, Foxp3+ regulatory T cells in human autoimmune disease. Nat Med 2011, 17(6): 673–675.

9. McClymont SA, Putnam AL, Lee MR, Esensten JH, Liu W, Hulme MA, et al. Plasticity of human regulatory T cells in healthy subjects and patients with type 1 diabetes. J Immunol 2011, 186(7): 3918–3926.

10. Lowther DE, Goods BA, Lucca LE, Lerner BA, Raddassi K, van Dijk D, et al. PD-1 marks dysfunctional regulatory T cells in malignant gliomas. JCI Insight 2016, 1(5).

11. Rubtsov YP, Rasmussen JP, Chi EY, Fontenot J, Castelli L, Ye X, et al. Regulatory T cell-derived interleukin-10 limits inflammation at environmental interfaces. Immunity 2008, 28(4): 546–558.

12. Laidlaw BJ, Cui W, Amezquita RA, Gray SM, Guan T, Lu Y, et al. Production of IL-10 by CD4(+) regulatory T cells during the resolution of infection promotes the maturation of memory CD8(+) T cells. Nat Immunol 2015, 16(8): 871–879.

13. Marson A, Housley WJ, Hafler DA. Genetic basis of autoimmunity. J Clin Invest 2015, 125(6): 2234–2241.

14. International Multiple Sclerosis Genetics C, Hafler DA, Compston A, Sawcer S, Lander ES, Daly MJ, et al. Risk alleles for multiple sclerosis identified by a genomewide study. N Engl J Med 2007, 357(9): 851–862.

15. International Multiple Sclerosis Genetics C, Nikolaos Patsopoulos, Sergio E. Baranzini, Adam Santaniello PS, Chris Cotsapas, Garrett Wong, Ashley H. Beecham, Tojo James, Joseph Replogle, Ioannis Vlachos, Cristin McCabe, Tune Pers, Aaron Brandes, Charles White, Brendan Keenan, Maria Cimpean, Phoebe Winn, Ioannis-Pavlos Panteliadis, Allison Robbins, Till F. M. Andlauer, Onigiusz Zarzycki, Benedicte Dubois, An Goris, Helle Bach Sondergaard, Finn Sellebjerg, Per Soelberg Sorensen, Henrik Ullum, Lise Wegner Thoerner, Janna Saarela, Isabelle Cournu-Rebeix, Vincent Damotte, Bertrand Fontaine, Lena Guillot-Noel, Mark Lathrop, Sandra Vukusik, Achim Berthele, Viola Biberacher, Dorothea Buck, Christiane Gasperi, Christiane Graetz, Verena Grummel, Bernhard Hemmer, Muni Hoshi, Benjamin Knier, Thomas Korn, Christina M Lill, Felix Luessi, Mark Muhlau, Frauke Zipp, Efthimios Dardiotis, Cristina Agliardi, Antonio Amoroso, Nadia Barizzone, Maria Donata Benedetti, Luisa Bernardinelli, Paola Cavalla, Ferdinando Clarelli, Giancarlo Comi, Daniele Cusi, Federica Esposito, Laura Ferre, Daniela Galimberti, Clara Guaschino, Maurizio A. Leone, Vittorio Martinelli, Lucia Moiola, Marco Salvetti, Melissa Sorosina, Domizia Vecchio, Andrea Zauli, Silvia Santoro, Miriam Zuccala, Julia Mescheriakova, Cornelia van Duijn, Steffan D. Bos, Elisabeth G. Celius, Anne Spurkland, Manuel Comabella, Xavier Montalban, Lars Alfredsson, Izaura L. Bomfim, David Gomez-Cabrero, Jan Hillert, Maja Jagodic, Magdalena Linden, Fredrik Piehl, Ilijas Jelcic, Roland Martin, Mireia Sospedra, Amie Baker, Maria Ban, Clive Hawkins, Pirro Hysi, Seema Kalra, Fredrik Karpe, Jyoti Khadake, Genevieve Lachance, Paul Molyneux, Matthew Neville, John Thorpe, Elizabeth Bradshaw, Stacy J. Caillier, Peter Calabresi, Bruce A. C. Cree, Anne Cross, Mary F. Davis, Paul de Bakker, Silvia Delgado, Marieme Dembele, Keith Edwards, Kate Fitzgerald, Irene Y. Frohlich, Pierre-Antoine Gourraud, Jonathan L. Haines, Hakon Hakonarson, Dorlan Kimbrough, Noriko Isobe, Ioanna Konidari, Ellen Lathi, Michelle H. Lee, Taibo Li, David An, Andrew Zimmer, Albert Lo, Lohith Madireddy, Clara P. Manrique, Mitja Mitrovic, Marta Olah, Ellis Patrick, Margaret A. Pericak-Vance, Laura Piccio, Cathy Schaefer, Howard Weiner, Kasper Lage, - ANZgene, - IIBDGC, - WTCCC2, Alastair Compston, David Hafler, Hanne F. Harbo, Stephen L. Hauser, Graeme Stewart, Sandra D’Alfonso, Georgios Hadjigeorgiou, Bruce Taylor, Lisa F. Barcellos, David Booth, Rogier Hintzen, Ingrid Kockum, Filippo Martinelli-Boneschi, Jacob L. McCauley, Jorge R. Oksenberg, Annette Oturai, Stephen Sawcer, Adrian J. Ivinson, Tomas Olsson, Philip L. De Jager. The Multiple Sclerosis Genomic Map: Role of peripheral immune cells and resident microglia in susceptibility. bioRxiv 2017, https://www.biorxiv.org/content/early/2017/07/13/143933.

16. Olsson T, Barcellos LF, Alfredsson L. Interactions between genetic, lifestyle and environmental risk factors for multiple sclerosis. Nat Rev Neurol 2017, 13(1): 25–36.

17. Kleinewietfeld M, Manzel A, Titze J, Kvakan H, Yosef N, Linker RA, et al. Sodium chloride drives autoimmune disease by the induction of pathogenic TH17 cells. Nature 2013, 496(7446): 518–522.

18. Wu C, Yosef N, Thalhamer T, Zhu C, Xiao S, Kishi Y, et al. Induction of pathogenic TH17 cells by inducible salt-sensing kinase SGK1. Nature 2013, 496(7446): 513–517.

19. Hernandez AL, Kitz A, Wu C, Lowther DE, Rodriguez DM, Vudattu N, et al. Sodium chloride inhibits the suppressive function of FOXP3+ regulatory T cells. J Clin Invest 2015, 125(11): 4212–4222.

20. Wei Y, Lu C, Chen J, Cui G, Wang L, Yu T, et al. High salt diet stimulates gut Th17 response and exacerbates TNBS-induced colitis in mice. Oncotarget 2017, 8(1): 70–82.

21. Tubbs AL, Liu B, Rogers TD, Sartor RB, Miao EA. Dietary Salt Exacerbates Experimental Colitis. J Immunol 2017, 199(3): 1051–1059.

22. Paling D, Solanky BS, Riemer F, Tozer DJ, Wheeler-Kingshott CA, Kapoor R, et al. Sodium accumulation is associated with disability and a progressive course in multiple sclerosis. Brain 2013, 136(Pt 7): 2305–2317.

23. Fitzgerald KC, Munger KL, Hartung HP, Freedman MS, Montalban X, Edan G, et al. Sodium intake and multiple sclerosis activity and progression in BENEFIT. Ann Neurol 2017, 82(1): 20–29.

24. Clevers H. Wnt/beta-catenin signaling in development and disease. Cell 2006, 127(3): 469–480.

25. Naito AT, Sumida T, Nomura S, Liu ML, Higo T, Nakagawa A, et al. Complement C1q activates canonical Wnt signaling and promotes aging-related phenotypes. Cell 2012, 149(6): 1298–1313.

26. Niehrs C. The complex world of WNT receptor signalling. Nat Rev Mol Cell Biol 2012, 13(12): 767–779.

27. Staal FJ, Luis TC, Tiemessen MM. WNT signalling in the immune system: WNT is spreading its wings. Nat Rev Immunol 2008, 8(8): 581–593.

28. Ding Y, Shen S, Lino AC, Curotto de Lafaille MA, Lafaille JJ. Beta-catenin stabilization extends regulatory T cell survival and induces anergy in nonregulatory T cells. Nat Med 2008, 14(2): 162–169.

29. van Loosdregt J, Fleskens V, Tiemessen MM, Mokry M, van Boxtel R, Meerding J, et al. Canonical Wnt signaling negatively modulates regulatory T cell function. Immunity 2013, 39(2): 298–310.

30. Keerthivasan S, Aghajani K, Dose M, Molinero L, Khan MW, Venkateswaran V, et al. beta-Catenin promotes colitis and colon cancer through imprinting of proinflammatory properties in T cells. Sci Transl Med 2014, 6(225): 225ra228.

31. Sorcini D, Bruscoli S, Frammartino T, Cimino M, Mazzon E, Galuppo M, et al. Wnt/beta-Catenin Signaling Induces Integrin alpha4beta1 in T Cells and Promotes a Progressive Neuroinflammatory Disease in Mice. J Immunol 2017, 199(9): 3031–3041.

32. Kitz A, de Marcken M, Gautron AS, Mitrovic M, Hafler DA, Dominguez-Villar M. AKT isoforms modulate Th1-like Treg generation and function in human autoimmune disease. EMBO Rep 2016, 17(8): 1169–1183.

33. Dias S, D’Amico A, Cretney E, Liao Y, Tellier J, Bruggeman C, et al. Effector Regulatory T Cell Differentiation and Immune Homeostasis Depend on the Transcription Factor Myb. Immunity 2017, 46(1): 78–91.

34. Kitagawa Y, Ohkura N, Kidani Y, Vandenbon A, Hirota K, Kawakami R, et al. Guidance of regulatory T cell development by Satb1-dependent super-enhancer establishment. Nat Immunol 2017, 18(2): 173–183.

35. Wu Y, Borde M, Heissmeyer V, Feuerer M, Lapan AD, Stroud JC, et al. FOXP3 controls regulatory T cell function through cooperation with NFAT. Cell 2006, 126(2): 375–387.

36. Pabbisetty SK, Rabacal W, Volanakis EJ, Parekh VV, Olivares-Villagomez D, Cendron D, et al. Peripheral tolerance can be modified by altering KLF2-regulated Treg migration. Proc Natl Acad Sci U S A 2016, 113(32): E4662–4670.

37. Lengfeld JE, Lutz SE, Smith JR, Diaconu C, Scott C, Kofman SB, et al. Endothelial Wnt/beta-catenin signaling reduces immune cell infiltration in multiple sclerosis. Proc Natl Acad Sci U S A 2017, 114(7): E1168–E1177.

38. Tawk M, Makoukji J, Belle M, Fonte C, Trousson A, Hawkins T, et al. Wnt/beta-catenin signaling is an essential and direct driver of myelin gene expression and myelinogenesis. J Neurosci 2011, 31(10): 3729–3742.

39. Staal FJ, van Noort M, Strous GJ, Clevers HC. Wnt signals are transmitted through N-terminally dephosphorylated beta-catenin. EMBO Rep 2002, 3(1): 63–68.

40. Duhen T, Duhen R, Lanzavecchia A, Sallusto F, Campbell DJ. Functionally distinct subsets of human FOXP3+ Treg cells that phenotypically mirror effector Th cells. Blood 2012, 119(19): 4430–4440.

41. Wing K, Onishi Y, Prieto-Martin P, Yamaguchi T, Miyara M, Fehervari Z, et al. CTLA-4 control over Foxp3+ regulatory T cell function. Science 2008, 322(5899): 271–275.

42. Harada N, Tamai Y, Ishikawa T, Sauer B, Takaku K, Oshima M, et al. Intestinal polyposis in mice with a dominant stable mutation of the beta-catenin gene. EMBO J 1999, 18(21): 5931–5942.

43. Sebastian M, Lopez-Ocasio M, Metidji A, Rieder SA, Shevach EM, Thornton AM. Helios Controls a Limited Subset of Regulatory T Cell Functions. J Immunol 2016, 196(1): 144–155.

44. Essers MA, de Vries-Smits LM, Barker N, Polderman PE, Burgering BM, Korswagen HC. Functional interaction between beta-catenin and FOXO in oxidative stress signaling. Science 2005, 308(5725): 1181–1184.

45. Okada K, Naito AT, Higo T, Nakagawa A, Shibamoto M, Sakai T, et al. Wnt/beta-Catenin Signaling Contributes to Skeletal Myopathy in Heart Failure via Direct Interaction With Forkhead Box O. Circ Heart Fail 2015, 8(4): 799–808.

46. Ouyang W, Liao W, Luo CT, Yin N, Huse M, Kim MV, et al. Novel Foxo1-dependent transcriptional programs control T(reg) cell function. Nature 2012, 491(7425): 554–559.

47. Dehner M, Hadjihannas M, Weiske J, Huber O, Behrens J. Wnt signaling inhibits Forkhead box O3a-induced transcription and apoptosis through up-regulation of serum-and glucocorticoid-inducible kinase 1. J Biol Chem 2008, 283(28): 19201–19210.

48. Wang R, Ferraris JD, Izumi Y, Dmitrieva N, Ramkissoon K, Wang G, et al. Global discovery of high-NaCl-induced changes of protein phosphorylation. Am J Physiol Cell Physiol 2014, 307(5): C442–454.

49. Irarrazabal CE, Burg MB, Ward SG, Ferraris JD. Phosphatidylinositol 3-kinase mediates activation of ATM by high NaCl and by ionizing radiation: Role in osmoprotective transcriptional regulation. Proc Natl Acad Sci U S A 2006, 103(23): 8882–8887.

50. Fang D, Hawke D, Zheng Y, Xia Y, Meisenhelder J, Nika H, et al. Phosphorylation of beta-catenin by AKT promotes beta-catenin transcriptional activity. J Biol Chem 2007, 282(15): 11221–11229.

51. McCubrey JA, Steelman LS, Bertrand FE, Davis NM, Abrams SL, Montalto G, et al. Multifaceted roles of GSK-3 and Wnt/beta-catenin in hematopoiesis and leukemogenesis: opportunities for therapeutic intervention. Leukemia 2014, 28(1): 15–33.

52. Fang X, Yu SX, Lu Y, Bast RC, Jr., Woodgett JR, Mills GB. Phosphorylation and inactivation of glycogen synthase kinase 3 by protein kinase A. Proc Natl Acad Sci U S A 2000, 97(22): 11960–11965.

53. Wehbi VL, Tasken K. Molecular Mechanisms for cAMP-Mediated Immunoregulation in T cells - Role of Anchored Protein Kinase A Signaling Units. Front Immunol 2016, 7: 222.

54. Lund RJ, Loytomaki M, Naumanen T, Dixon C, Chen Z, Ahlfors H, et al. Genome-wide identification of novel genes involved in early Th1 and Th2 cell differentiation. J Immunol 2007, 178(6): 3648–3660.

55. Boniface K, Bak-Jensen KS, Li Y, Blumenschein WM, McGeachy MJ, McClanahan TK, et al. Prostaglandin E2 regulates Th17 cell differentiation and function through cyclic AMP and EP2/EP4 receptor signaling. J Exp Med 2009, 206(3): 535–548.

56. Kofler DM, Marson A, Dominguez-Villar M, Xiao S, Kuchroo VK, Hafler DA. Decreased RORC-dependent silencing of prostaglandin receptor EP2 induces autoimmune Th17 cells. J Clin Invest 2014, 124(6): 2513–2522.

57. Li X, Murray F, Koide N, Goldstone J, Dann SM, Chen J, et al. Divergent requirement for Galphas and cAMP in the differentiation and inflammatory profile of distinct mouse Th subsets. J Clin Invest 2012, 122(3): 963–973.

58. Sreeramkumar V, Fresno M, Cuesta N. Prostaglandin E2 and T cells: friends or foes? Immunol Cell Biol 2012, 90(6): 579–586.

59. Yao C, Hirata T, Soontrapa K, Ma X, Takemori H, Narumiya S. Prostaglandin E(2) promotes Th1 differentiation via synergistic amplification of IL-12 signalling by cAMP and PI3-kinase. Nat Commun 2013, 4: 1685.

60. Shin H, Kwack MH, Shin SH, Oh JW, Kang BM, Kim AA, et al. Identification of transcriptional targets of Wnt/beta-catenin signaling in dermal papilla cells of human scalp hair follicles: EP2 is a novel transcriptional target of Wnt3a. J Dermatol Sci 2010, 58(2): 91–96.

61. Castellone MD, Teramoto H, Williams BO, Druey KM, Gutkind JS. Prostaglandin E2 promotes colon cancer cell growth through a Gs-axin-beta-catenin signaling axis. Science 2005, 310(5753): 1504–1510.

62. Notani D, Gottimukkala KP, Jayani RS, Limaye AS, Damle MV, Mehta S, et al. Global regulator SATB1 recruits beta-catenin and regulates T(H)2 differentiation in Wnt-dependent manner. PLoS Biol 2010, 8(1): e1000296.

63. Pappalardo JL, Hafler DA. The Human Functional Genomics Project: Understanding Generation of Diversity. Cell 2016, 167(4): 894–896.

64. Wilck N, Matus MG, Kearney SM, Olesen SW, Forslund K, Bartolomaeus H, et al. Salt-responsive gut commensal modulates TH17 axis and disease. Nature 2017.

65. Vojdani A. A Potential Link between Environmental Triggers and Autoimmunity. Autoimmune Dis 2014, 2014: 437231.

66. Shao J, Jung C, Liu C, Sheng H. Prostaglandin E2 Stimulates the beta-catenin/T cell factor-dependent transcription in colon cancer. J Biol Chem 2005, 280(28): 26565–26572.

67. Mahic M, Yaqub S, Johansson CC, Tasken K, Aandahl EM. FOXP3+CD4+CD25+ adaptive regulatory T cells express cyclooxygenase-2 and suppress effector T cells by a prostaglandin E2-dependent mechanism. J Immunol 2006, 177(1): 246–254.

68. Kihara Y, Matsushita T, Kita Y, Uematsu S, Akira S, Kira J, et al. Targeted lipidomics reveals mPGES-1-PGE2 as a therapeutic target for multiple sclerosis. Proc Natl Acad Sci U S A 2009, 106(51): 21807–21812.

